# Aging-associated alternative splicing programs conserved between human and mouse tissues

**DOI:** 10.64898/2025.12.30.697080

**Authors:** Keren Isaev, David A. Knowles

## Abstract

Alternative splicing is a key gene regulatory process that diversifies the proteome and controls gene dosage. Previous studies have detected aging-associated splicing changes across various tissues. However, their use of bulk RNA-seq obfuscates the impacted cell-types and may confound cell-type proportion changes with cell-intrinsic ones. We present a framework that first assembles and maps alternative splicing events in appropriate single cell RNA-seq data (scRNA-seq), then applies LeafletFA, a probabilistic model that discovers coordinated splicing programs (SPs) without requiring prior knowledge of cell types or clinical information, such as age. Applying this framework to over 200,000 cells from mouse and human Smart-seq2 multi-tissue atlases, we discovered global and cell type specific aging-associated SPs. In mice, integrating SPs with gene expression significantly improved age prediction in 46 of 76 tissue-cell types tested. We identified the RNA-binding protein *Snrnp70* as a key upstream mediator driving the shift from youthful to aged SPs. To test evolutionary conservation of these patterns, we employed transfer learning to map the murine splicing dictionary onto human transcriptomes. This revealed a conserved “youth” program (SP4) that is maintained in quiescent endothelial tissues but lost in high-turnover organs. We identified a conserved, age-dependent alternative 5’ splice site usage in the splicing factor *SRSF5* as a molecular marker of this program in both species. Our work establishes alternative splicing as a coordinated, evolutionarily conserved dimension of cellular aging and provides LeafletFA as an open-source tool for single-cell splicing analysis.

## Introduction

Aging is a complex biological process characterized by a progressive loss of transcriptional and protein homeostasis, increasing the risk of chronic conditions such as neurodegenerative disorders, cardiovascular disease, and cancer (1). Although changes in gene expression during aging have been extensively studied, including at single-cell resolution, the contribution of alternative splicing (AS) to this process has only been explored using bulk RNA-seq. AS enables single genes to generate multiple RNA and protein variants through selective exon inclusion and the use of alternative splice sites (SS), allowing cells to modulate RNA and protein function, localization, and stability in response to developmental and environmental signals (2). AS can also impact gene dosage by altering mRNA stability, translation efficiency, or inducing nonsense-mediated decay. How AS homeostasis contributes to cellular aging and how its dysregulation drives age-related decline remains unclear (3).

Expression-centric single-cell RNA sequencing studies have revealed inter-cell-type divergent and asynchronous aging trajectories(4). In the mouse brain, aging-associated gene expression changes are highly cell-type-specific and can even be opposing; for example, cholesterol metabolism genes are downregulated in oligodendrocytes while being upregulated in certain neurons (5). In transdifferentiated neurons, aging is associated with the mislocalization of essential spliceosome components from the nucleus to the cytoplasm leading to widespread AS and cellular stress (6). It remains unclear, however, whether these changes reflect an increase in functional complexity (7) or a more widespread loss of splicing fidelity with age (8; 9; 10). Because molecular aging patterns are often highly cell-type-specific, single-cell approaches are uniquely suited to address this question. Full-length scRNA-seq methods like Smart-seq2 are required for such an analysis, as they capture internal splicing events missed by the more widespread 3’ tagging approaches such as 10x. Such approaches have revealed a layer of cell-type-specific splicing information, complementary to gene expression, in datasets like the Tabula Muris and mouse cortex (11; 12; 13; 14). However, these studies were limited in scope, focusing on cell-type differences in AS within single tissue. A systematic, cross-tissue, single-cell analysis of aging-associated AS has yet to be performed. We fill this gap by uniform processing, integration, and factor modeling of large scale Smart-seq2 atlases: Tabula Muris Senis (15), Tabula Sapiens (16), and Allen Brain datasets (17).

Comprehensive, scalable and unbiased characterization of AS in these atlas scale datasets is not possible with existing software. For instance, tools like SpliZ require predefined cell-type annotations and provide only gene-level scores (18; 19), while methods such as PsiX are restricted to annotated exons and rely on gene expression to define cell similarity (20). These constraints prevent the discovery of splicing programs (SPs), i.e. coordinated sets of splicing events that collectively contribute to a cell state (21; 22; 23; 24; 25). This is analogous to well established gene expression programs, in which a biological state, such as inflammation, is defined by the coordinated activation of a module of genes (26). An aging SP would therefore represent a module of co-varying splicing events that reflect an age-associated cellular state. By focusing on individual splicing events or being tethered to gene expression, current tools cannot uncover these system-level splicing regulatory programs across the cellular diversity of large aging atlases.

To address these challenges, we developed LeafletFA. Our framework first creates a large-scale single-cell splicing resource by systematically processing existing Smart-seq2 atlases. It then employs a novel Bayesian factor model to discover latent SPs. By applying LeafletFA to over 200,000 cells and nuclei from mouse and human datasets spanning multiple ages, we provide the first comprehensive analysis of how AS variation relates to cellular aging. Our approach reveals age-associated splicing signatures that are both distinct from and complementary to gene expression changes, offering new insights into the molecular complexity of aging.

## Results

### Detection of Single-Cell Alternative Splicing Variation Across Species

To build a comprehensive catalog of alternative splicing (AS) events, we compiled splice junction data from large-scale Smart-seq2-based atlases. These short-read, full-length transcript coverage methods are well-suited for robust splice junction identification (**Fig. 1A**). Our integrated dataset encompasses over 200,000 cells and nuclei from mouse (Tabula Muris Senis, Allen Brain Atlas) and human (Tabula Sapiens, Allen Brain Atlas) across diverse tissues and a wide range of ages. We processed the mouse and human datasets by extracting splice junctions from split reads using RegTools (27). To ensure complete and high-resolution splice site (SS) annotation, we leveraged a bulk PacBio long-read RNA-sequencing (LRS) derived GTF file (28; 29). This LRS-derived annotation provided an exhaustive catalog of known and novel full-length transcript isoforms and their associated SSs in both species, allowing us to more accurately map and classify the diverse junctions observed in the short-read data. Groups of overlapping splice junctions representing distinct AS events were then aggregated into Alternative Transcript Structure Events (ATSEs), analogous to LeafCutter clusters (30), using our <u>ATSEmapper module.</u> ATSEs reflect AS and alternative first/last exon choices and capture transcript structure diversity (**Fig. 1B**). We obtained 113,995 junctions in 32,567 ATSEs (12,307 genes) in mouse (**Fig. 1C-D**), and 160,251 junctions in 41,489 ATSEs (13,668 genes) in human (**Supp. Fig. 1A**).

**Fig. 1:**
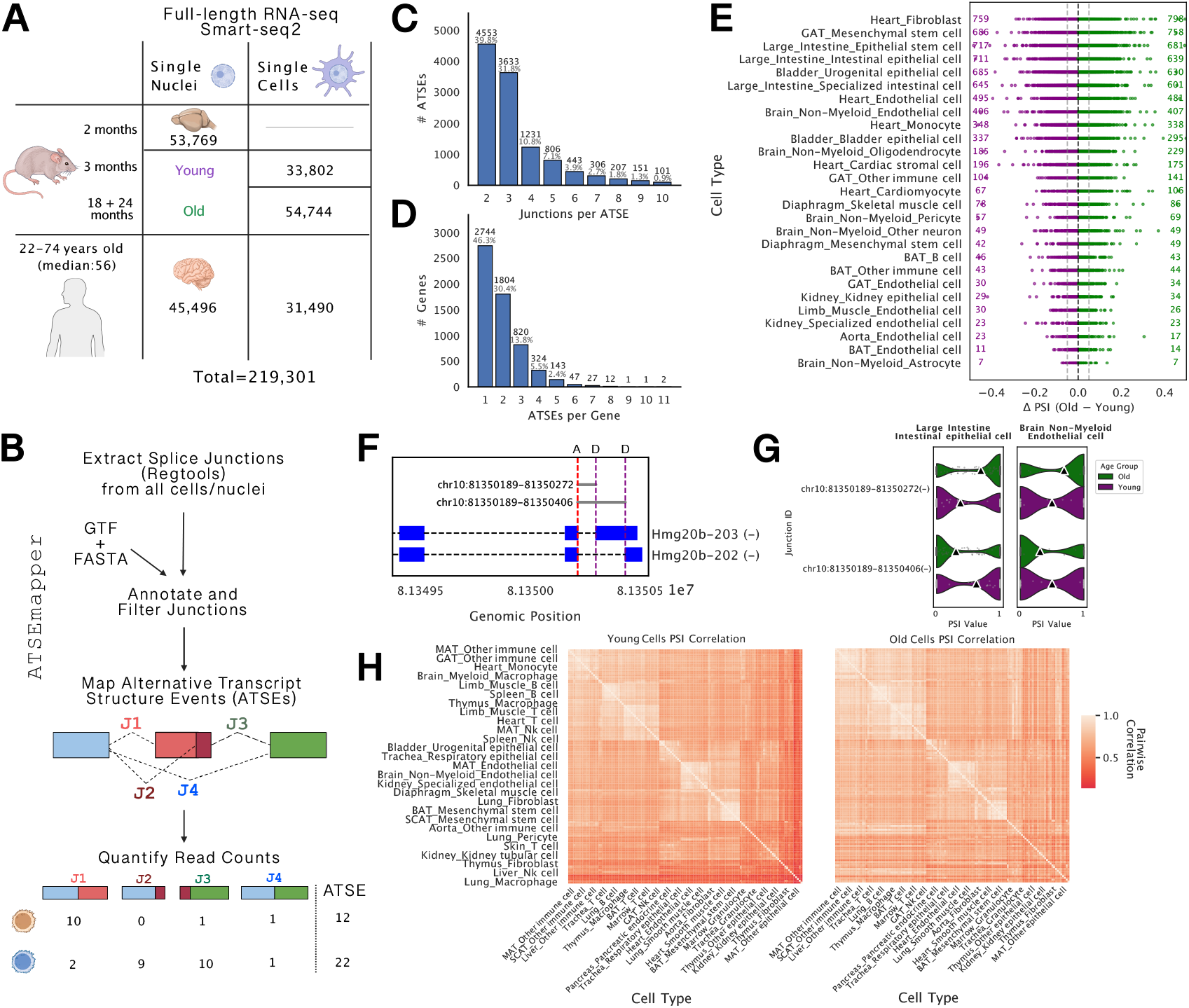
Overview of data processing and alternative splicing event quantification. **(A)** Summary of single-cell and single-nuclei RNA-seq datasets (Smart-seq2) included in the study, categorized by age group in mouse and human. **(B)** The ATSEmapper quantification pipeline. Splice junctions are extracted using RegTools and a reference GTF/FASTA file, filtered, and then mapped to define Alternative Transcript Structure Events (ATSEs). The output includes quantified read counts for all junctions (J1-J4 shown as an example) comprising the ATSE in each cell/nuclei. **(C)** Distribution of the number of splice junctions per ATSE in mouse. **(D)** Distribution of the number of ATSEs per gene in mouse. **(E)** Distribution of age-associated alternative splicing changes (*Δ*PSI) across tissue-cell type pairs in mouse cells (3-24 months). Each dot represents the top splice junction within an ATSE, selected by having the highest absolute *Δ*PSI (PSI_Old_ − PSI_Young_) in that cell type. The flanking numbers indicate the count of junctions with absolute *Δ*PSI > 0.05 (dashed lines), representing shifts toward lower (purple) or higher (green) PSI in old cells. **(F)** Example ATSE in the *Hmg20b* gene (chr10). The splice graph schematic shows two annotated isoforms (Hmg20b-203 and Hmg20b-202) and highlights the aging associated junctions corresponding to two Alternative Donor (D) sites (A: acceptor). **(G)** Violin plots showing the distribution of observed PSI values for the splice junctions in (F), comparing Young (purple) vs. Old (green) cells in specific cell types (Large Intestine Intestinal epithelial cells and Brain Non-Myeloid Endothelial cells). **(H)** Pairwise correlation matrices of PSI values between broad cell types, separated by age group (Left: Young Cells; Right: Old Cells). The matrices were generated by correlating mean PSI values per cell type across all junctions. Clustering was performed only on the Young Cells matrix, and the resulting row/column order was applied to the Old Cells matrix to allow for direct comparison.**Abbreviations:** ATSE, Alternative Transcript Structure Event; PSI, Percent Spliced In; BAT, Brown adipose tissue; GAT, Gonadal adipose tissue; MAT, Mesenteric adipose tissue; NK, Natural killer; SCAT, Subcutaneous adipose tissue.

As expected, the less-annotated splice junctions were detected in significantly fewer cells, highlighting a challenge in testing the splicing fidelity hypothesis: the very junctions that might represent age-related errors are captured too sparsely (**Supp. Fig. 1B**). Nevertheless, to probe for age-related AS changes, we performed a preliminary analysis by calculating the direct difference in observed junction usage (*Δ*PSI) between old (18 and 24 months) and young cells (3 months) in mice, where percent spliced in (PSI) is the ratio of junction count to total ATSE count for each junction within each cell. We identified 11,875 junctions with age-associated shifts (|*Δ*PSI| > 0.05) in at least one tissue-cell type pair, with the highest number observed in heart fibroblasts (**Fig. 1E**, **Supp. Fig. 1D**).

A representative example is the chromatin factor *Hmg20b*, which undergoes a consistent age-dependent shift in alternative donor site usage (**Fig. 1F**). While youthful cells favor the distal donor site (*Hmg20b-203*), older cells increasingly utilize the proximal site (*Hmg20b-202*). This shift is reproducible across distinct lineages, including intestinal epithelial cells in the large intestine and non-myeloid endothelial cells in the brain (**Fig. 1G**).

Beyond analyzing individual cell types, we assessed global pairwise shifts in PSI. By correlating observed junction PSI across cell type pairs, we found that older cells are more similar to each other in their AS profiles compared to younger cells (**Fig. 1H**). This increased similarity suggests global erosion of cell-type specific AS during aging. However, this approach is limited by the sparse nature of single-cell data. The ability to detect a change was strongly correlated with how frequently a junction was observed across cells (Spearman’s *ρ* = 0.7), suggesting the analysis lacks the statistical power to uncover subtle but biologically meaningful changes in most junctions. Moreover, the identification of SPs is essential for understanding how cells coordinate post-transcriptional regulation (21; 23). This systems-level perspective is particularly crucial for aging analysis, where dysregulation may manifest as aberrant activation of entire SPs rather than random errors in individual junctions. These challenges underscore the need for a probabilistic framework that can handle data sparsity to reliably identify differential splicing.

### LeafletFA Decomposes Alternative Splicing Variation into Splicing Programs

To address the sparsity and complexity of single-cell splicing data, we developed LeafletFA, a probabilistic Beta-Dirichlet factor model. LeafletFA decomposes the sparse cell-by-junction count matrix into a set of interpretable, continuous, latent SPs (**Fig. 2A**). The model represents each cell’s splicing profile as a weighted combination of these programs (via a priori Dirichlet-distributed factor weights, *Φ*) and defines each program by a vector of usage ratios for each junction (a priori Beta-distributed usage ratios, *Ψ*). We fit LeafletFA using Stochastic Variational Inference via the Pyro probabilistic programming language (31), ensuring LeafletFA is highly scalable and computationally efficient for atlas-level datasets. Model parameters and methodology are detailed in **[1]**

**Fig. 2:**
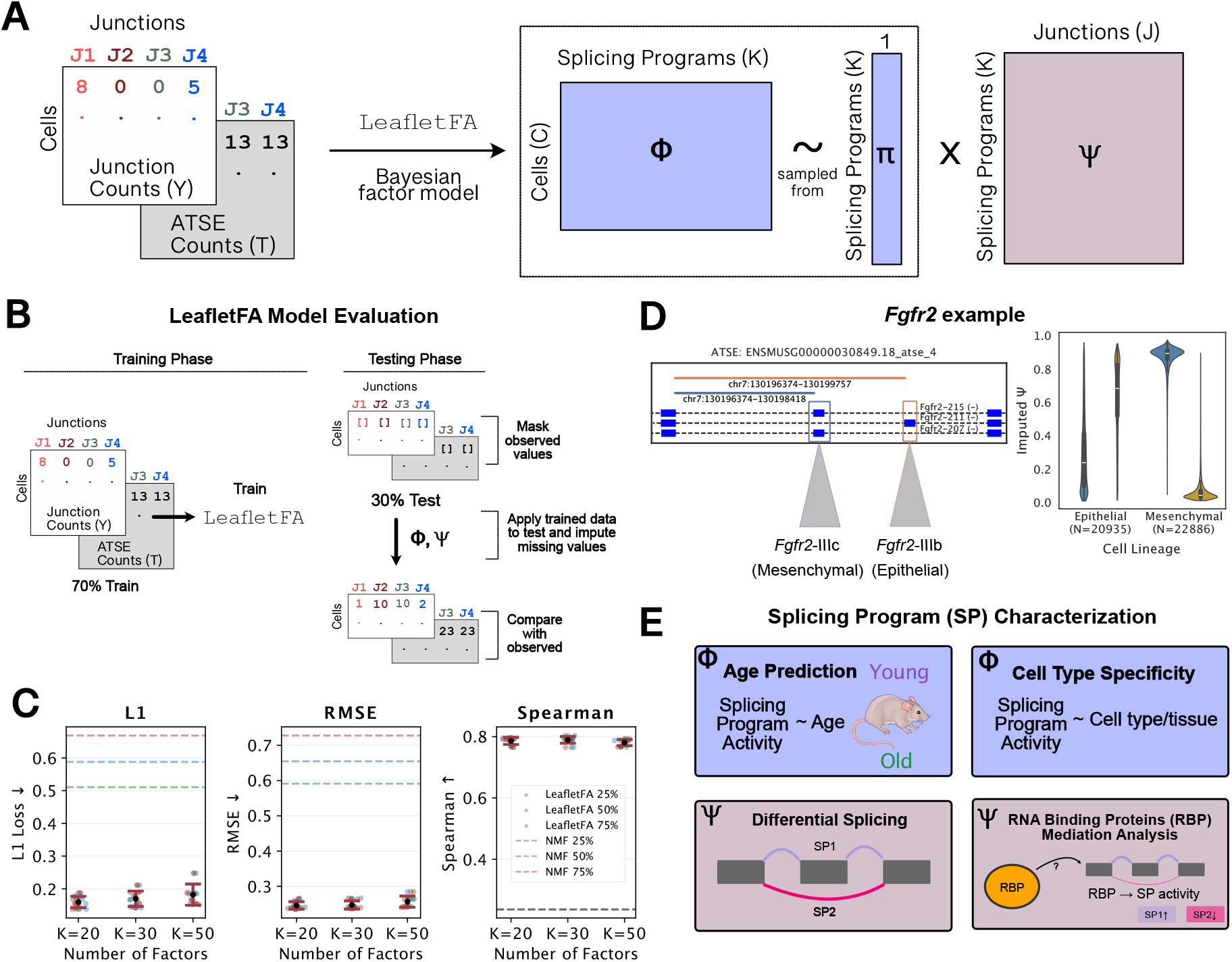
Architecture and Validation of the LeafletFA Bayesian Factor Model for Decomposing Single-Cell Splicing Variation. **(A)** Schematic representation of the LeafletFA (Factor Analysis) Bayesian factor model. The model decomposes the sparse Junction Counts matrix (*Y*) for *J* junctions across *C* cells into a Cell Factor Matrix (**Φ**) and a SP Matrix (**Ψ**) to identify *K* latent SPs. These SPs capture co-occurring alternative splicing events (ATSEs). **(B)** Overview of the Leaflet-FA Model Evaluation strategy to assess imputation accuracy. The Junction Counts (*Y*) for a subset of cells are partially masked. The model is trained on the remaining data (70% Train and unmasked test data) and then used to impute the masked junction counts. Model performance is evaluated by comparing the imputed values (*Ψ, Φ*) against the original observed values. **(C)** Evaluation of LeafletFA’s imputation accuracy compared to NMF across various masking percentages (25%, 50%, 75%) and different numbers of factors (*K* = 20, *K* = 30, *K* = 50). Performance is measured by RMSE ↓ (Root Mean Square Error), L1 Loss ↓, and Spearman ↑ (Spearman correlation coefficient). **(D)** Example of the identified *Fgfr*2 ATSE in mouse cells, which illustrates the imputation process for an event linked to Cell Lineage (Mesenchymal *Fgfr*2 − *IIIc* vs. Epithelial *Fggr*2− *IIIb* isoforms). **(E)** Applications of LeafletFA for SP Characterization. The latent SP activities (*Φ*) are biologically characterized by associating them with various covariates: Cell Type Specificity (e.g., Mesenchymal vs. Epithelial lineages), Age Prediction (linking activity to Young vs. Old age groups), and Mediation Analysis to identify regulatory relationships between RNA Binding Proteins (RBP), their expression, and SP activity.

We rigorously validated LeafletFA using both simulated and real-world benchmarks. First, using simulated data where the ground truth structure was known, we confirmed the model’s accuracy in recovering latent SPs and cellular activities, observing superior performance compared to Non-negative Matrix Factorization (NMF) (**Supp. Fig. 2**). Second, we assessed imputation capabilities on the integrated single-cell mouse splicing atlas using a masking strategy. 70% of junction-cell pairs are used as training data, and the remaining 30% as test data (**Fig. 2B**). LeafletFA significantly outperformed NMF in recovering these masked values across multiple metrics, validation percentages (25%, 50%, 75%), and latent factor dimensions (*K* = 20, 30, 50) (**Fig. 2C**).

LeafletFA’s ability to recover biologically relevant splicing patterns in real data is exemplified by the *Fgfr2* gene, a canonical example of mutually exclusive exon usage driven by epithelial-mesenchymal transition (32). LeafletFA correctly imputed the binary switch between the epithelial isoform (*Fgfr2-IIIb*) and the mesenchymal isoform (*Fgfr2-IIIc*) (**Fig. 2D**). By modeling the entire dataset collectively, LeafletFA leverages information across all cells and junctions to resolve splicing decisions that would be ambiguous in sparse raw data.

Having established the model’s accuracy on simulated benchmarks and its robustness in imputing sparse data, we defined a workflow to interpret the discovered latent factors. The SPs identified by LeafletFA can be biologically characterized by correlating their cellular activity (*Φ*) with cell lineage and biological age, and by identifying regulatory RNA-binding proteins (RBPs) through mediation analysis (**Fig. 2E**). This framework allows us to move beyond individual junction metrics and detect coordinated, organism-wide post-transcriptional programs.

### LeafletFA Reveals Interpretable Splicing Programs Associated with Mouse Aging

To dissect AS programs in the aging mouse, we applied LeafletFA to a comprehensive atlas of 142,315 single cells and nuclei spanning **159 unique tissue-cell type pairs**. We initialized the model with *K* = 20 factors (SPs), which upon training showed varying levels of global utilization (*π*). Ordering SPs by their *π* value, SP1-SP5 exhibited the largest contributions to overall splicing variation (total percentage of variance explained=0.98) across cells (**Fig. 3A**). The SPs have quite low correlation within one another, with the exception of some small clusters, e.g. SP5-7 and SP3/8/10, suggesting they capture distinct AS signatures (**Fig. 3B**).

**Fig. 3:**
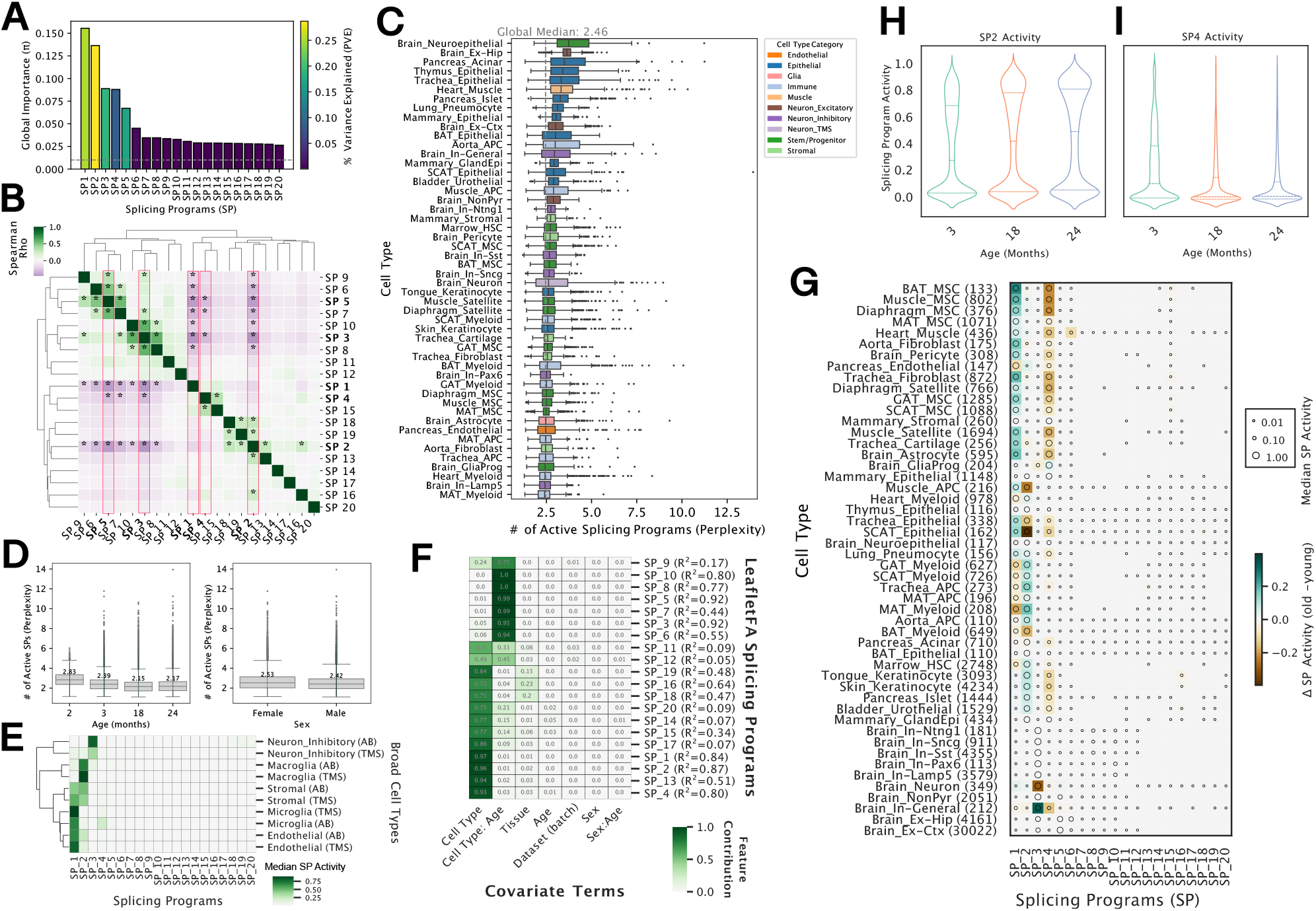
LeafletFA reveals interpretable splicing programs and aging signatures in mouse cells. **(A)** Global usage parameter (*π*) values for each SP (SP1-SP20) defined by LeafletFA. Bars are colored by the percentage of variance explained (PVE). **(B)** Pairwise Spearman correlation matrix of SP activity (*ϕ*), organized by hierarchical clustering to reveal correlated SPs. SP1-SP5 are highlighted due to high PVE. Stars indicate a p-value < 0.05 in the Spearman correlation. **(C)** Distribution of SP perplexity (effective number of active SPs) by cell type. Only the top 50 cell types (out of 159 unique tissue-cell type pairs) with the highest median perplexity are shown. The global median perplexity is 2.46. Full perplexity results for all cell types are provided in **Table S1. (D)** Distribution of SP perplexity across age groups (2m, 3m, 18m, 24m) and sex (Female, Male). **(E)** Agreement of SP usage profiles across datasets (Allen Brain vs. Tabula Muris Senis) shown by hierarchical clustering of median SP activity (*ϕ*). **(F)** Feature contribution heatmap showing the proportion of variance (*R*^2^) in each SP explained by covariates, derived from ANOVA. Rows are ordered by hierarchical clustering. **(G)** Age-associated changes in SP activity for the top 50 cell types (out of 159 total). The heatmap displays the median difference in activity (*Δ*SP Activity: Old - Young). **Brown** indicates higher activity in young mice (negative *Δ*), while **Teal/Blue** indicates higher activity in old mice (positive *Δ*). Full differential activity results are provided in **Table S2. (H, I)** Temporal dynamics of SP2 (**H**) and SP4 (**I**) activity across three age points (3, 18, and 24 months), shown as violin plots. **Abbreviations:** APC, Antigen-presenting cell; BAT, Brown adipose tissue; Brain_Ex-Hip, Excitatory neurons of the hippocampus; Brain_In, Inhibitory neurons; GAT, Gonadal adipose tissue; HSC, Hematopoietic stem cell; MAT, Mesenteric adipose tissue; MSC, Mesenchymal stem cell; NonPyr, Non-pyramidal neurons; SCAT, Subcutaneous adipose tissue.

We quantified SP complexity as the effective number of active SPs per cell (i.e., perplexity, see Methods). While the global median perplexity was 2.46, this metric varied markedly across tissue-cell types (**Fig. 3C**; the 50 types with the highest complexity are shown). Excitatory neurons and epithelial cells exhibited the highest complexity (median perplexity of 3.09 and 2.61, respectively), whereas microglia and macroglia (astrocytes and oligodendrocytes) cells displayed the lowest (median of 1.60 and 1.79). Notably, the number of active programs appeared to decline with age (**Fig. 3D**, left), although this trend may be partially driven by the enrichment of high-complexity neurons in the 2-month-old samples. Finally, we observed a minor sex bias, with female cells retaining slightly higher complexity than male cells (median perplexity of 2.53 vs. 2.42; **Fig. 3D**, right).

To assess the robustness of the SPs, we examined their agreement across the distinct datasets used (Allen Brain nuclei and Tabula Muris Senis cells). Hierarchical clustering of median SP activity demonstrated that similar cell types cluster together regardless of the source dataset, confirming that LeafletFA recovers biological signals rather than batch effects (**Fig. 3E**).

To further analyze the variation in SP activities, we performed a global analysis of variance (ANOVA) (**Fig. 3F**). For 13 programs (including SP1, SP2, and SP4), the specific tissue-cell type accounts for the majority of the observed variance. The remaining seven programs, such as SP3, SP5, and SP10, are primarily driven by the interaction between tissue-cell type and age. Notably, even when tissue-cell type defines baseline activity, these SPs could still capture significant age-related shifts within those contexts. Most importantly, technical covariates such as dataset batch explained negligible variance, typically less than 1% across nearly all programs.

To resolve context-specific aging signals, we examined changes in SP activity within the top 50 cell types from **Fig. 3C** between old and young mice (**Fig. 3G**). This revealed a striking directionality in specific programs. SP1 and SP2 tend to increase with age (indicated by teal shifts), while SP4 activity consistently decreases (indicated by brown shifts). These trends were further corroborated by analyzing the temporal dynamics of SP2 and SP4, which show progressive shifts in activity distribution from 3 to 24 months (**Fig. 3H, I**). Unlike these systemic aging programs, other SPs displayed restricted age effects (e.g., SP6 in muscle), highlighting the model’s ability to deconvolve complex AS landscapes into both systemic and cell-type-specific aging signatures.

To investigate the interplay between post-transcriptional and transcriptional regulation during aging, we correlated the 20 LeafletFA SPs (*ϕ*) with 20 latent gene expression factors (*Z*) learned by linear scVI (**Fig. 3I**). Although significant, the correlations were generally moderate (|*ρ*| < 0.5), suggesting that splicing captures cell-state information that is complementary to, rather than redundant with, gene expression. We found that the aging-correlated SPs identified in **Fig. 3G** map to distinct, and often opposing, transcriptional landscapes. SP1 and SP2, which both increase with age but in distinct lineages, align with divergent gene signatures. SP1 (aging stroma) positively correlates with inflammatory and stromal factors like Z16, Z6, and Z1. In contrast, SP2 (aging immune/epithelial) exhibits an inverse relationship with these same factors. Conversely, the youth-associated program SP4 exhibits a ubiquitous profile, appearing across diverse lineages (**Fig. 3G**). This broad utility is reflected in its complex transcriptional coupling: SP4 tracks with neuronal identity via a negative correlation with Z_8_, yet simultaneously captures non-neuronal signals via a positive correlation with Z_10_ highlighting that SP4 is not lineage-restricted, but likely represents a pan-tissue regulatory program of “youthful” splicing that is active across both neuronal and non-neuronal cell contexts.

### Integration of Splicing and Expression Reveals Complementary Aging Signatures

Motivated by these distinct signatures and by previous successes of DNA methylation and protein clocks (33; 34; 35; 36), we assessed the predictive power of each modality using ridge regression to predict chronological age (**Fig. 4A**). For this analysis, we subset the data to 76 tissue-cell type pairs (*N* = 80, 557 cells) with *n* ≥ 50 cells in both young and old groups. Across these cell types, gene expression alone was often a stronger age predictor than splicing alone, consistent with previous reports linking transcriptional changes to aging (37; 38). However, incorporating SPs alongside gene expression significantly improved performance: a nested-model ANOVA comparing the combined model (splicing + expression) against the expression-only model revealed significant performance gains in 46 of the 76 (61%) tissue-cell types analyzed (FDR < 0.05). This broad improvement underscores that SPs provide independent age-related information not fully captured by gene expression changes alone.

**Fig. 4:**
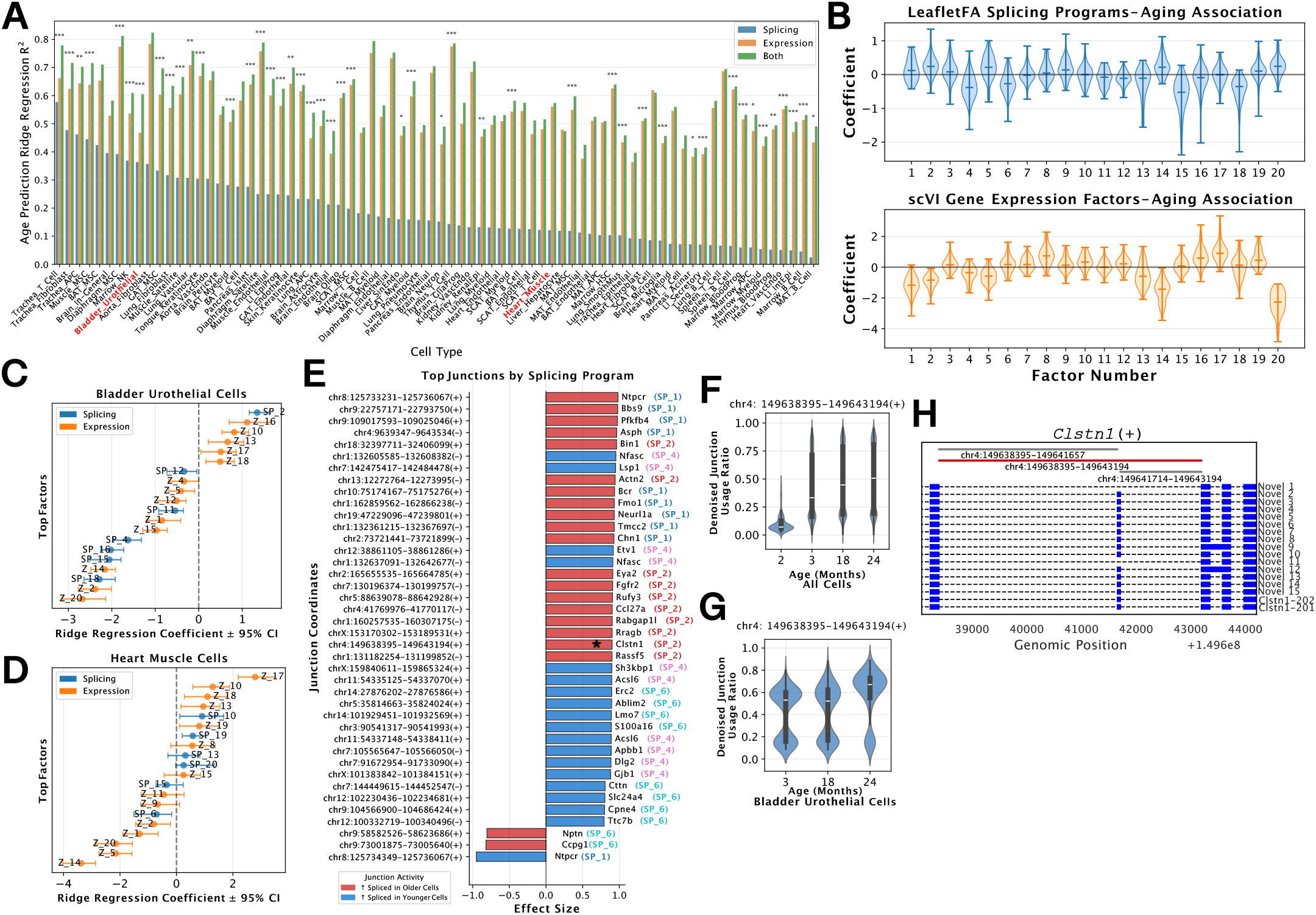
Integration of splicing and expression features reveals cell type-specific aging signatures. **(A)** Age prediction performance (Ridge regression *R*^2^) across cell types comparing three feature sets: SPs alone (blue), gene expression factors alone (orange), and combined features (green). For this in-depth analysis of aging, the dataset was subset to 76 tissue-cell types (N=80,557 cells) containing at least 50 cells in both young and old age groups. Asterisks indicate a significant improvement (*** FDR < 0.001) in the combined model compared to gene expression alone. **(B)** Distribution of aging association coefficients across cell types from the combined Ridge regression models in (A). Top panel shows coefficients for each SP while the bottom panel shows similar analysis for gene expression factors. **(C-D)** Cell type-specific aging signatures. Forest plots showing Ridge regression coefficients (*±* 95% CI using bootstrapping) for top contributing features in **(C)** Bladder Urothelial Cells and **(D)** Heart Muscle Cells. **(E)** Top splice junction markers for aging-correlated SPs. Bar plots show the top junction effect sizes for programs such as SP1, SP2, SP4 and SP6. Red bars indicate junctions with increased splicing in older cells, while blue bars indicate increased splicing in younger cells. **(F-G)** Temporal dynamics of the top SP2-associated splice junction in *Clstn1* (chr4:149638395-149643194(+)). Violin plots show the denoised junction usage ratio (from LeafletFA) across age time points (2, 3, 18, and 24 months all cells (**(F)**)) and just Bladder Urothelial Cells (**(G)**). **(H)** Genomic context of the *Clstn* junction shown in (G). Schematic illustrates the reference and novel transcripts containing the splice junctions in the ATSE.

SP1 and SP2 generally displayed positive coefficients (aging-associated) and SP4 trended negative (youth-associated) when fit jointly with gene expression scVI factors. However, these averages masked significant heterogeneity across tissues (**Fig. 4B**). This suggests that rather than acting as a uniform global clock, aging involves a mixture of global and context-specific effects, where the magnitude and direction of splicing shifts are modulated by the specific requirements of each cell type. Notably, even SPs with lower global utilization frequently obtained strong regression coefficients (SP15 and SP18 for example), demonstrating that they capture age-related variance distinct from the transcriptional background. These context-specific dynamics are evident when examining individual cell types. For example, in bladder urothelial cells (**Fig. 4C**), SP2 is the strongest positive predictor of age out of all features (transcriptional or splicing), while SP4 is a robust negative predictor alongside several other SPs. In heart muscle cells (**Fig. 4D**), expression factors like Z_17_ (linked to *Grik1* and *Il1rapl2*, **Fig. 3I**) were dominant positive predictors; yet, splicing features still contributed significantly, with SP6 as a strong negative predictor. SPs provided this power despite inherent mathematical constraints. While scVI Z factors are unconstrained latent variables, SPs are compositionally constrained (summing to 1 per cell). That they robustly capture aging variance despite this constraint highlights the biological importance of AS shifts.

To resolve the molecular targets of the SPs, we next identified the top splice junction markers for aging-associated SPs using effect sizes estimated from LeafletFA’s variational parameters (**Fig. 4E**, Methods). For a given SP, a positive effect size indicates that the junction is used more frequently in that SP relative to all other programs. One of the top hits for the aging-associated SP2 was a splice junction in *Clstn1* (chr4:149638395-149643194(+)), a gene typically expressed in neurons. As expected usage of this junction increases with age broadly across cells (Fig.4F) as well as in specific cell types, e.g. bladder urothelial cells (Fig.4G). Mapping this junction to the genomic context reveals it corresponds to a novel splice exon-skipping event captured by long-read annotations (Fig.4H), demonstrating that single-cell splicing analysis can uncover potential aging-related splice events missed by standard references.

### Mediation Analysis Identifies RNA-Binding Proteins as Drivers of Aging-Associated Splicing Programs

To define the specific splicing events governed (markers) by each SP, we performed differential splicing (DS) analysis on the SP junction usage rates (*Ψ*), leveraging the variational posterior to estimate uncertainty. The SPs learn distinct regulatory footprints containing thousands of significant junctions (**Fig. 5A**). Interestingly, the aging-associated programs (SP1, SP2) and the youth-associated program (SP4) ranked among the highest in junction count and contained a notable number of novel or unannotated splice junctions. This suggests that aging likely involves widespread transcriptome remodeling mediated by changes in SPs, rather than isolated stochastic errors.

**Fig. 5:**
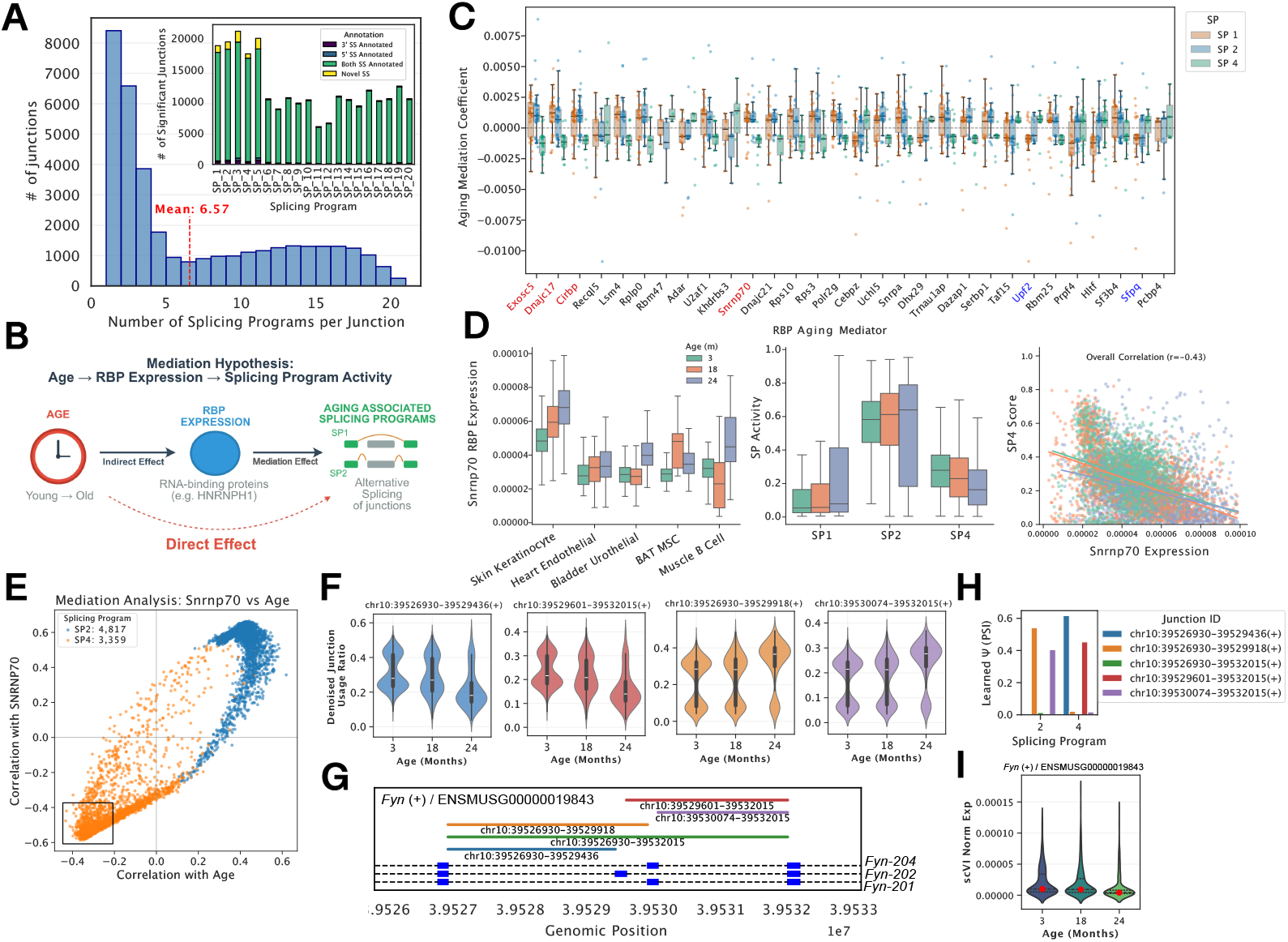
Global analysis of splicing program dynamics and their RBP mediators during aging. **(A)** Global landscape of SP events. Histogram showing the distribution of junction specificity, defined as the number of SPs (out of 20 total) in which each junction is significant. The majority of junctions are specific to a single program, while a subset shows pleiotropic activity across multiple programs. Inset shows the total count of significant junctions captured per program by annotation status. **(B)** Schematic of the mediation hypothesis: chronological age drives changes in RBP expression (the mediator), which in turn regulates the activity of specific SPs (outcome). **(C)** Systematic identification of aging mediators. The aging mediation coefficient for the top 30 RNA-binding protein (RBP) candidates across cell types is shown for SPs 1, 2 and 4. Genes highlighted in red exhibit consistently positive coefficients for aging-associated SP1 and SP2, and negative coefficients for the youth-associated SP4 while those highlighted in blue show the opposite trend. **(D)** *Snrnp70* expression and SP dynamics. **Left:** Expression of *Snrnp70* consistently increases with age (3m to 24m) across multiple tissues (skin, heart, bladder, BAT, muscle). **Middle:** Activity of SPs shifts with age; the aging-associated program (SP2) increases, while the youth-associated program (SP4) declines. **Right:** Scatter plot showing negative correlation (*r* = − 0.43) between *Snrnp70* expression and the youth splicing score (SP4) in single cells. **(E)** Global mediation analysis in bladder urothelial cells. Scatter plot showing the correlation of individual splice junction usage (markers of SP2 and SP4) with age (x-axis) versus their correlation with *Snrnp70* expression (y-axis). The concordance reveals that junctions lost with age (markers of SP4-youth program, bottom-left) are repressed by *Snrnp70*, while those gained (markers of SP2-aging program, top-right) are upregulated. **(F)** Violin plots showing the distribution of denoised, per-cell junction usage ratios (imputed PSI) for four specific *Fyn* splice junctions across age groups in bladder urothelial cells. Colors indicating specific junctions are consistent across panels (F)-(H). **(G)** Genomic track of the *Fyn* gene (ENSMUSG00000019843) illustrating the genomic coordinates and connectivity of the differentially regulated AS junctions analyzed in (F). **(H)** Bar plot showing the learned inclusion levels (*Ψ*) for five junctions within the *Fyn* locus for SP 2 (aging) and 4 (youth). These values represent the global splicing dictionary learned by LeafletFA for each SP. **(I)**Violin plot showing the distribution of *scVI* normalized gene expression for *Fyn* across the 3, 18, and 24-month age groups. **Abbreviations:** BAT, Brown adipose tissue.

We next asked whether SP activity is driven by specific RNA Binding Proteins (RBPs). We constructed a mediation model, hypothesizing that age alters the expression of specific RBPs (the mediators), which in turn drives the activity of aging SPs (SP1,SP2 and SP4, **Fig. 5B**). We calculated an Aging Mediation Coefficient for 269 RBPs to quantify their role in propagating aging signals. RBPs often exhibit high co-expression (**Supp. Fig. 5A**), implying that observed mediation effects may capture redundant regulatory modules rather than unique drivers. Furthermore, these mediation effects displayed significant heterogeneity across cell types (**Fig. 5C**, **Supp. Fig. 5B**).

Despite this heterogeneity, we identified a set of RBPs that exhibit opposing mediation effects on youth versus aging programs. Specifically, RBPs with positive mediation effects in aging programs (SP1 and SP2) consistently displayed negative mediation effects for the youth program (SP4) (**Fig. 5C**). Prominent examples of this trend include *Exosc5, Dnajc17, Cirbp* and *Snrnp70*. Higher expression of these RBPs is associated with an overall aging phenotype defined by the reduced activity of SP4. Conversely, RBPs such as *Upf2* and *Sfpq* exhibited the inverse pattern: high positive mediation effects for SP4 and negative coefficients for SP1 and SP2 (**Supp. Fig. 5C**). These patterns suggest that the aging splicing phenotype does not arise from a single isolated pathway, but rather from a coordinated shift in the expression of multiple core splicing factors that collectively favor ‘aging’ programs over ‘youth’ programs.

We identified the U1 snRNP component *Snrnp70* as a top-ranked mediator of this transition. *Snrnp70* expression increases with age across diverse tissues, including skin, heart, bladder, and muscle (**Fig. 5D**, left). This age-dependent upregulation correlates with the activation of the aging SPs and the concurrent repression of the youth program in these cell types (**Fig. 5D**, middle). Single-cell correlation analysis confirmed a negative relationship (*r* = − 0.43) between *Snrnp70* expression and the SP4 Youth Score (**Fig. 5D**, right).

To investigate this relationship further, we focused on bladder urothelial cells, a context where we previously established that SP2 and SP4 are key coefficients in the aging model even when controlling for gene expression (**Fig. 4C**). To determine if *Snrnp70* acts as a direct regulator of these programs, we extracted differentially spliced junctions for the aging (SP2) and youth (SP4) programs (**Fig. 5A**). We then correlated the imputed PSI values of these program-specific junctions with both chronological age and *Snrnp70* expression. We observed a global concordance (**Fig. 5E**): junctions belonging to the aging-associated SP2 consistently showed positive correlations with both age and *Snrnp70*, while junctions from the youth program (SP4) displayed coincident negative correlations. The observed coupling suggests that changes in *Snrnp70* activity impact these junctions, making its age-dependent upregulation a major contributor to the splicing shifts that define the aging transcriptome.

A prime example of this regulation is observed in the cytoskeletal regulator *Fyn*. In bladder urothelial cells, aging drives complex splicing shifts characterized by the coordinated loss of youth-enriched junctions (markers of SP4) and the concurrent gain of aging-enriched junctions (markers of SP2) (**Fig. 5F**, **Supp. Fig. 5D**). This transition encompasses mutually exclusive exon skipping events; specifically, the blue and red junctions dominant in younger cells (enriched in SP4 and found in *Fyn-202*) are lost with age, while the orange and purple junctions (representing *Fyn-204* and *Fyn-201*) increase in usage. Genomic visualization of the *Fyn* locus illustrates the connectivity of these differentially regulated junctions (**Fig. 5G**). While these splicing patterns shift dramatically, total *Fyn* expression decreases only slightly across the age groups (**Fig. 5I**). This highlights how age-dependent upregulation of mediators like *Snrnp70* may impact the AS landscape through largely independent of changes in total gene abundance.

### Cross-Species Transfer Learning Reveals Conserved Aging-associated Splicing Programs

To determine if the age-associated SPs identified in mice are evolutionarily conserved, we utilized transfer learning to map mouse splicing signatures onto human transcriptomes. We curated a human Smart-seq2 dataset from the Tabula Sapiens (*N* = 31, 490 cells) and Allen Brain (*N* = 45, 496 nuclei) atlases. Given the smaller sample size and genetic and environmental diversity present in the human cohort, we opted not to train a new human model from scratch. Instead, we tested if the “dictionary” of SPs learned in the mouse (*Ψ*_*M*_) could explain splicing variation in human cells (**Fig.6A**). By fixing the program definitions to those learned in the mouse and restricting the feature space to 13,510 strictly orthologous splice junctions (Methods), we inferred SP activities (*Φ*_*H*_) for human cells.

**Fig. 6:**
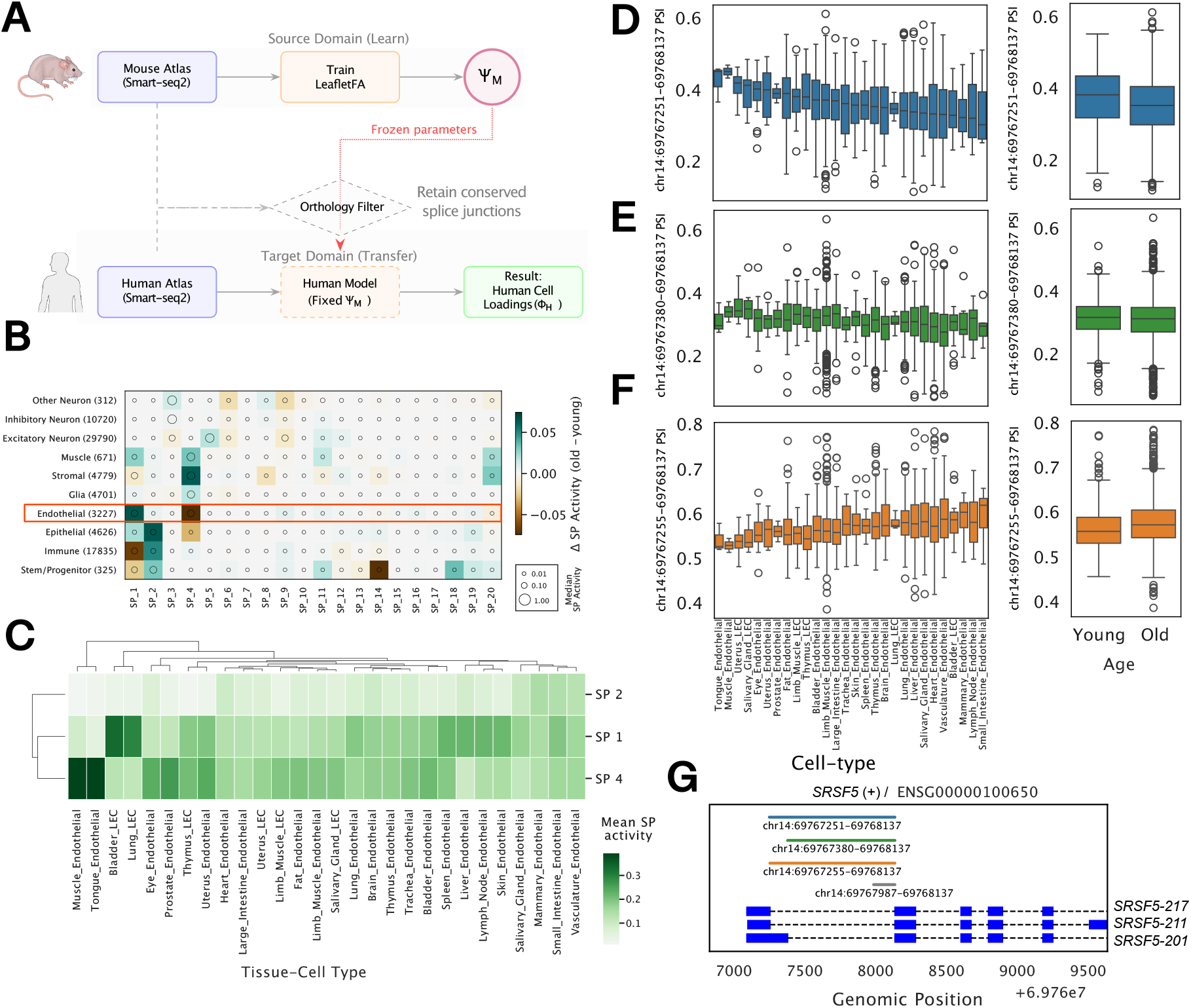
Transfer learning reveals conserved age-associated splicing programs in human endothelial cells. Schematic of the LeafletFA transfer learning workflow. SP dictionaries (*Ψ*_*M*_) obtained from the trained mouse LeafletFA model (**Figures 3–5**) are transferred to the human Smart-seq2 dataset. Human cell loadings (*Φ*_*H*_) are inferred using only orthologous splice junctions (*N* = 13, 510) detected in both species. **(B)** Dot plot showing the change in SP activity (Old minus Young) across broad human cell types. Dot size represents median SP activity, and color indicates the magnitude of age-associated change. The endothelial lineage (highlighted) exhibits distinct age-dependent program shifts. **(C)** Hierarchical clustering of human endothelial subtypes based on the mean activity of three mouse-defined SPs (SP 1, SP 2, and SP 4). **(D-F)** Box plots showing imputed Percent Spliced In (PSI) for three junctions in the human *SRSF5* locus across diverse endothelial tissue-cell types (left) and young versus old cohorts (right). Panel (D) features the youth-associated SP4 marker (decreasing with age), while panel (F) features the SP1-associated aging marker located four nucleotides upstream (increasing with age). **(G)** Gene structure and isoform visualization for human *SRSF5*. The colors of the splice junction tracks correspond to the ones analyzed in (D-F), highlighting the usage of alternative 5’ SSs. **Abbreviations:** LEC, Lymphatic endothelial cell;

Despite the constraints of cross-species transfer, the model successfully recovered broad human cell identities solely based on conserved splicing patterns (**Supp. Fig. 6A**). Inferred SP activity is largely conserved between mouse and human, particularly for the primary aging programs: SP1, SP2, and SP4. In both species, SP4 is broadly active across various lineages, mirroring its murine distribution with high median activity in stromal, glial, muscle, and endothelial populations (**Fig.6B**). This cell-type concordance extends to the aging-associated SP1 and SP2, which show moderate to high baseline activity in endothelial and epithelial lineages across both datasets. Beyond these broad lineages, SP5 demonstrates a highly specific conserved activity pattern restricted primarily to excitatory neurons in both species (**Fig.6B**). Notably, immune populations exhibit a comparable profile characterized by higher SP1 activity in younger cells that transitions toward increased SP2 activity in older cells.

While the SP-based cellular identities are consistent, the conservation of aging trajectories, represented by *ΔSP* activity, varies across broad cell types. The strongest and most consistently conserved aging signal emerged within the diverse endothelial lineage (**Fig.6B**). The age-associated transition within these endothelial populations is highly conserved across species and is defined by a significant induction of SP1 accompanied by a reduction in the youthful SP4 (**Fig.6B-C**, **Supp. Fig. 6B**). Program usage is stratified by the physiological demands of the tissue rather than species-specific factors. Endothelial cells from structurally stable and quiescent vascular beds, such as those in muscle, maintain high baseline activity of the youthful SP4 in both mice and humans (**Fig.6C**). In contrast, endothelial cells from high-turnover filtration organs, such as the liver, exhibit a higher baseline activity of the age-associated SP1 program. This global concordance across human cells and the original mouse atlas suggests that the shift from SP4 to SP1 could reflect a universal biological response to physiological turnover and cumulative tissue stress.

To identify conserved aging markers, we screened the 13,510 orthologous junctions and subset to the 5,710 that served as significant markers for either SP1 or SP4 in the mouse. We validated these markers in human endothelial cells (*n* = 3, 227) and found that 2,811 (49%) were significantly age-correlated in the human cohort, where significance was defined as an absolute Spearman’s *ρ* > 0.05 and an adjusted *p*-value < 0.05. Notably, 2,336 of these significant junctions (83%) maintained the same directionality of change observed in the mouse, confirming a high degree of evolutionary conservation in structural aging.

An example of this is an ATSE in *SRSF5* (Serine/Arginine-Rich Splicing Factor 5). The imputed activity (PSI) of two junctions within this locus, as learned by LeafletFA, is significantly correlated with age in human endothelial cells. Specifically, the most upstream junction (blue; chr14:69767255-69768137) acts as a marker of the aging-associated SP1, while the junction situated four nucleotides downstream (chr14:69767251-69768137) is a marker of the youthful SP4, mirroring the regulatory architecture found in mouse cells. This ATSE involves an alternative 5’ SS and transcriptional start site (**Fig.6G**). In both species, the alternative SSs differ by exactly four nucleotides (**Supp. Fig. 6C**). These results demonstrate that splicing changes during aging are not random, but rather an evolutionarily conserved shift in post-transcriptional programs.

## Discussion

We developed LeafletFA, a scalable Bayesian factor model for analyzing AS at single-cell resolution, and created the first comprehensive atlas of splicing variation across mouse and human cells. By processing over 200,000 cells from multiple Smart-seq2 datasets, we demonstrated that AS provides a complementary layer of cellular identity that enhances our understanding of aging beyond gene expression alone.

Our analysis revealed several key insights. First, while SPs showed overall modest correlations with gene expression programs, the relationship between transcription and splicing is complex, involving co-transcriptional regulation and bidirectional influences (39; 40; 41; 42). This coupling occurs through dynamic spatial, physical, and temporal organizational properties, with splicing kinetics playing a crucial role in determining splicing outcomes (43; 44). Our latent factor approach captures the net result of these complex interactions, revealing that while splicing and expression are coupled processes, they still contribute distinct information about cellular states that cannot be fully predicted from one another.

Second, we identified global and context dependent aging signatures in AS. SP1, SP2 and SP4 in emerged as the primary aging-associated programs across both mouse and human. The cell type-specific variation in how strongly these programs correlate with age aligns with recent findings that different tissues and cell types age at different rates (45; 38). The cell type-specific nature of these aging programs may reflect differential expression of RBPs and splicing regulators across cell types, leading to distinct post-transcriptional responses to aging.

Our findings should be interpreted in light of emerging evidence about the population-specific nature of aging biomarkers. Recent work has shown that inflammaging, commonly considered a hallmark of aging, may not be universal across human populations (46). The inflammatory aging signature appears to be largely a product of industrialized lifestyles rather than an inherent biological aging process. This raises important questions about the generalizability of splicing-based aging signatures we observe, which are derived primarily from laboratory mice and individuals from industrialized populations. Future work should examine whether the aging SPs we identify are similarly influenced by environmental and lifestyle factors.

Several limitations merit consideration. The integration of single cells and single nuclei data remains challenging due to fundamental differences in RNA capture, with nuclear RNA’s inclusion of intronic reads potentially capturing different biological processes (47). Additionally, distinguishing functional splicing variation from unproductive splicing or technical artifacts remains difficult, as many detected events may represent splicing noise rather than regulated programs (48). The correlations we observe between splicing factors and gene expression could also partially reflect NMD-induced expression differences (49).

Our analysis does not capture intron retention events, which represent a significant component of age-related transcriptome deterioration. Recent work has shown that intron retention increases broadly with age across human tissues, contributing to a progressive dilution of functional mRNAs with non-functional splicing isoforms (10). While methods like FRASER have been developed to detect aberrant splicing including intron retention from RNA-seq data (50), our current approach focuses on alternative SS and exon skipping events. This limitation is particularly relevant given that intron retention may double the number of detectable aberrant splicing events and has been implicated in pathogenic mechanisms. Future iterations of LeafletFA will incorporate intron retention analysis to provide a more complete picture of age-related splicing deterioration.

Our current study analyzes splicing and expression separately before integration. Future developments should explore joint modeling frameworks, similar to MultiVI (51; 52; 53), that could capture the regulatory relationships between transcription and splicing more directly. Such approaches could better distinguish between coordinated and independent regulation of these processes. Additionally, our model treats age as a linear variable, and more flexible approaches are needed to capture the potentially non-linear aging trajectories that likely differ across cell types.

Future work should explore several key directions. Recent advances in deep single-cell transcriptomics have revealed remarkable cell-specific isoform diversity even among pan-neuronal genes (54), suggesting that neuronal SPs may be particularly complex and informative. Second, applying LeafletFA to emerging technologies like single-cell long-read sequencing (55) would enable analysis of full-length isoforms and coordination between multiple splicing events within genes. Third, scaling to larger datasets using combinatorial indexing approaches(56; 57) could reveal rare cell type-specific SPs.

In conclusion, LeafletFA provides a foundation for incorporating AS into single-cell atlases, revealing that splicing represents a crucial but underexplored dimension of cellular identity and aging. As single-cell technologies continue to evolve, integrating splicing analysis will be essential for a complete understanding of cellular heterogeneity in health and disease.

## Methods

### Single-Cell Smart-seq2 Data Processing

We processed publicly available datasets to generate inputs for the LeafletFA model training and evaluation (**Fig. 1A**). For mouse data, the genome reference GENCODE vM19 (GRCm38) was used for all alignments and annotations. For human data, GENCODE v45 (GRCh38) was to capture annotated splice junctions. Four primary data sources were utilized to cover a wide range of tissues and species:

1. **Tabula Muris Senis (TMS):** Single-cell Smart-seq2 BAM files from the Tabula Muris Senis consortium (15) were obtained from the Chan Zuckerberg Biohub AWS S3 bucket: s3://czb-tabula-muris-senis/ Processed count matrices for total gene expression were used for gene expression analysis.
2. **Allen Brain Atlas - Mouse:** Smart-seq2 based single-nuclei RNA-seq data (ssv4) from the Allen Institute for Brain Science (58) were obtained from the NCBI Gene Expression Omnibus (GEO accession: GSE185862) and the Allen Brain Map portal: https://portal.brain-map.org/atlases-and-data/rnaseq/mouse-whole-cortex-and-hippocampus-smart-seq Processed count matrices, specifically exon-only counts or adjusted intron and exon counts, were used for gene expression analysis.
3. **Tabula Sapiens:** Single-cell Smart-seq2 data from the Tabula Sapiens consortium (16; 59) were obtained from: https://tabula-sapiens-portal.ds.czbiohub.org/ Brain tissue samples were excluded to avoid overlap with the Allen Brain human data.
4. **Allen Brain Atlas - Human:** Human brain single-nuclei Smart-seq2 data from the Allen Institute were obtained from the Allen Brain Map portal:https://portal.brain-map.org/atlases-and-data/rnaseq/human-multiple-cortical-areas-smart-seq

### Gene Expression Normalization and Integration

To enable the joint analysis of single-cell and single-nuclei datasets, we implemented a multi-step normalization and integration pipeline. Following best practices for Smart-seq2 data analysis (60), we normalized raw gene counts to account for transcript length bias by dividing each gene’s counts by its mean transcript length and then multiplying by the overall median transcript length across all genes. For the Allen Brain nuclei data, which contains intronic reads, we applied the same normalization to intronic counts using mean intron lengths. After length normalization, counts underwent library size normalization and a log1p transformation.

Quality control filtering was applied to remove low-quality cells and potential outliers. We calculated batch-wise robust statistics using median and median absolute deviation (MAD) for gene count and total count metrics. Cells were filtered based on standardized scores (z-scores) calculated from these robust statistics, retaining only cells within ±3 standard deviations of the batch-specific median. Additional filters included removal of cells with >10% ribosomal RNA content and cells lacking cell type annotations. We also removed long non-coding RNAs (lncRNAs) from both the gene expression and splicing data objects.

To make the nuclear expression profiles from the Allen Brain data comparable to whole-cell measurements, we combined the length-normalized exonic and intronic counts and trained linear regression models on cell types common to both datasets. These models learned the relationship between nuclear and whole-cell expression, and the derived coefficients were used to adjust all nuclei expression values.

For subsequent analyses, we focused on the 2,000 most highly variable genes and applied dimensionality reduction approaches for gene expression analysis. We employed Linear scVI (61) using the length-normalized raw integer counts as input, with dataset origin (single-cell vs. single-nuclei) specified as a batch covariate to account for technical differences between profiling methods. scVI analyses were performed with a dimensionality of *K* = 20.

### Alternative Splicing Quantification and ATSE Mapping

Splice junctions were extracted from BAM files using regtools (27) and processed using our AT-SEmapper pipeline (https://github.com/daklab/LeafletFA-utils/tree/main/leafletfa_utils/atsemapper). Each identified splice junction was validated using species-specific reference genome FASTA files to confirm the presence of canonical SS motifs (GT-AG, GC-AG, or AT-AC). GTF annotation files were used to classify each SS as either annotated (matching known exon boundaries) or novel, allowing us to retain high-confidence novel splicing events while filtering out potential alignment artifacts.

A splice graph is constructed for each gene where nodes represent unique SS and edges represent the observed junctions connecting them. An Alternative Transcript Structure Event (ATSE), analogous to an “intron cluster” in LeafCutter (30), is defined as a set of two or more junctions that are interconnected through common SS, forming a single connected component within the gene’s splice graph. ATSEs capture various forms of alternative splicing including cassette exons, alternative 5’/3’ SS, and alternative first/last exon usage (**Fig. 1B**).

Junction coordinates were grouped into ATSEs within each species separately, with mouse and human datasets processed independently to maintain species-specific SS coordinates and annotations. Junctions were filtered according to read and cell count thresholds (minimum 10 reads and 5 cells per junction, with minimum intron lengths of 50 bp and maximum of 500,000 bp). The number of reads supporting each junction in each ATSE in each cell/nucleus were aggregated into sparse AnnData(62) objects for downstream analysis.

**Table 1:**
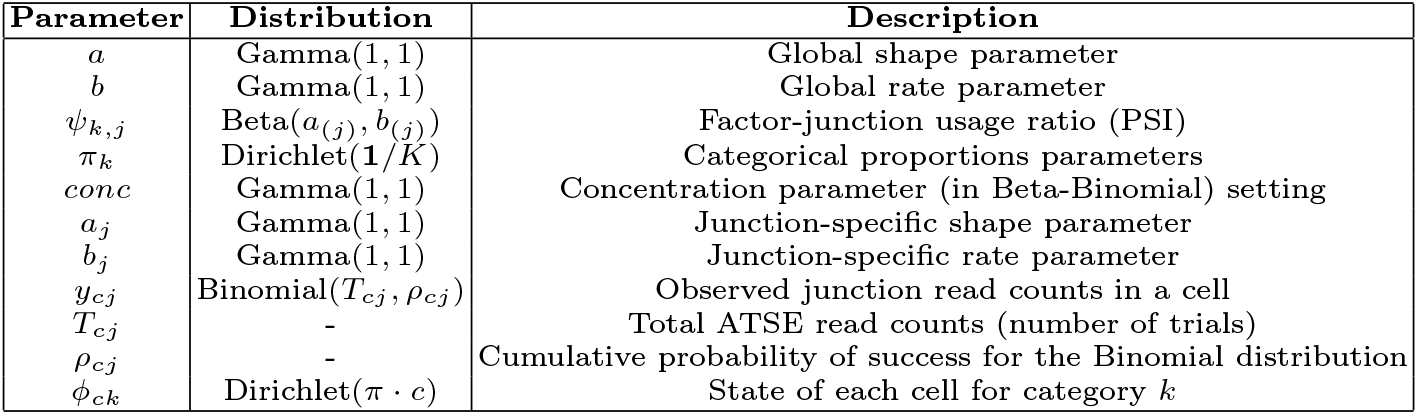
LeafletFA Model Parameters.

### LeafletFA Beta-Dirichlet Factor Model

LeafletFA models observed junction read counts *y*_*cj*_ conditional on total ATSE read counts *T*_*cj*_ for each cell *c* and junction *j*. We model each cell as a weighted combination *ϕ*_*c*:_ of *K* latent factors, each of which has a usage ratio *ψ*_*kj*_ for each junction,

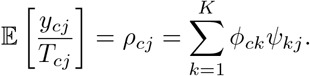

The prior for *ϕ* is given by the hierarchy,

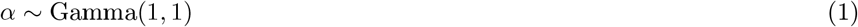

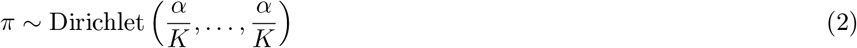

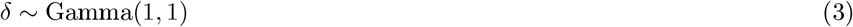

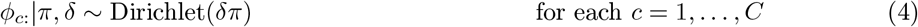

where *C* is the number of cells. We explored two different configurations for the prior on *ψ* but ultimately utilized the junction-specific prior configuration in our final LeafletFA model:

1. Global prior:

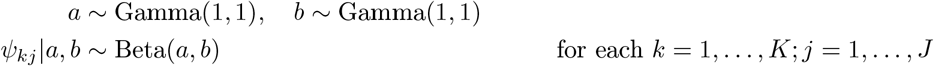

where *J* is the number of junctions.
2. Junction-specific priors:

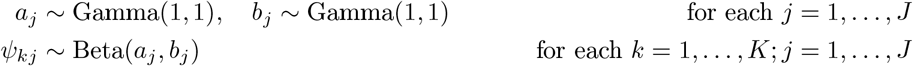

We considered two different likelihood models and implement the Beta-Binomial likelihood in the final model:
  - **Beta-binomial**:

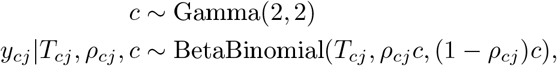

where *c* is a “concentration” parameter (inverse overdispersion). The beta-binomial represents additional noise in scenarios with high technical variability.
  - **Binomial:**

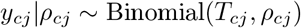

which we note is equivalent to the Beta-binomial when *c* → ∞.

### Multi-pass Mini-batch Training and Global Parameter Update

To enable scalable training on the large ~ 200,000 cell atlas, we implemented a multi-pass mini-batch training strategy optimized for GPU acceleration. The model was trained over 10 full passes of the data, with cells processed in mini-batches of 5,000.

In this approach, the global parameters *Ψ* (latent junction usage ratios) and *Π* (factor proportions) are treated as global variables that evolve continuously throughout the training process across all passes.

1. Batch Training: In each batch, LeafletFA trains the local parameters *Φ* (cell factor activities) specific to the 5,000 cells in that batch, while simultaneously refining the global parameters *Ψ* and *Π* using the Evidence Lower Bound (ELBO) objective.
2. Global Update and Re-initialization: After each mini-batch finishes training, the refined estimates of the global parameters (*Ψ, Π*) are extracted and stored. These refined parameters are then used to initialize the model for the *next* batch, ensuring that the model leverages the accumulated information from all previously processed cells.
3. Learning Rate Decay: A dynamic learning rate schedule was used to promote convergence, applying a decay factor of 0.95 to the base learning rate at the start of each new data pass. The first pass was trained for 50 epochs, while subsequent passes were trained for 20 epochs to conserve computational resources while benefiting from continuous global parameter refinement.

This strategy ensures all cells contribute to the final global factor structure (*Ψ* and *Π*) while efficiently utilizing GPU memory and computational resources.

### Model Fitting and Initialization Using Waypoints

We fit LeafletFA using Stochastic Variational Inference (SVI) within the Pyro probabilistic programming framework (31), optimizing the Evidence Lower Bound (ELBO).

Two initialization approaches were considered: random and waypoint-based. For the former the model is fit three times, and the one with the best ELBO was selected. We use waypoint-based initialization, inspired by Palantir (63), to reduce run-to-run variability due to optimization getting trapped in different local minima, and to improve reproducibility and potentially overall fit. This strategy is applied differently depending on the training context:

1. Initial Waypoint (First Batch): For the first mini-batch processed, the number of waypoints is set to the number of factors *K*. The waypoints are chosen to capture major subspaces of variation in the observed read count ratios, 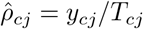. Given the sparsity in single-cell data, we first mean-centered non-missing values to reduce total expression influence. We extracted ten PCs from these adjusted junction usage ratios using sparse singular value decomposition (SVD). We use max-min sampling in the PC space to select *K* diverse cells as waypoints. The initial *ψ* for factor *k* is set to the average non-missing junction usage ratio across the 50 nearest neighbors (to reduce noise) of waypoint *k*, for each *k*.
2. Sequential Initialization (Subsequent Batches): For all subsequent mini-batches, the model is initialized using the global *Ψ* and *Π* estimates accumulated from the preceding batch, as detailed in **Section 1**. This continuous update strategy leverages the learning across all cells seen so far, ensuring efficient training.

### Bayesian Factor-Specific Differential Splicing Analysis

To identify splice junctions that exhibit significant differential usage across latent factors, we use a Bayesian posterior sampling-based approach inspired by scVI’s differential expression framework. Given the inferred latent variables from LeafletFA, we estimate factor-specific differential splicing by comparing the contribution of a given factor *k* to splice junction usage relative to the combined effect of the remaining factors.

#### Posterior Sampling and Effect Size Estimation

Since our approach is approximately Bayesian, we estimate differential splicing probabilistically by drawing posterior samples from the variational approximation. The default number of posterior samples is 500. Let *ψ*_*s,kj*_ denote the posterior sample *s* of *ψ*_*kj*_, and *ϕ*_*s,ck*_ posterior sample *s* for the factor assignment weights *ϕ*_*ck*_. For each sample, the factor-specific junction usage is,

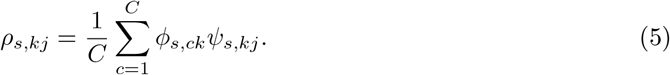

To estimate the overall effect of factor *k* relative to all others, we compute,

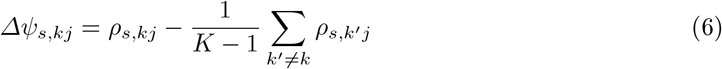

ensuring that the difference is computed against all other factors combined. Since *ψ*_*s,kj*_ and *ϕ*_*s,ck*_ are sampled from the posterior distribution, this formulation allows us to quantify uncertainty in differential splicing estimates.

#### Effect Size and Probabilistic Estimation of Differential Splicing

Rather than relying on a single effect size estimate, we use posterior sampling to empirically estimate the probability that a junction is differentially spliced. We define the overall effect size as,

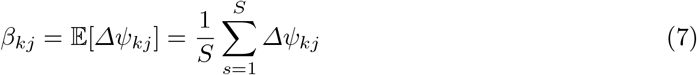

i.e., the mean factor-specific differential splicing effect across posterior samples. Additionally, we define the probability that a junction is differentially spliced as,

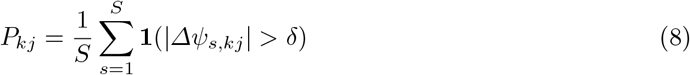

where *P*_*kj*_ is the fraction of posterior samples where the effect size exceeds the threshold *δ*. The default threshold *δ* is set to 0.1, but can be adjusted based on the variability of the data set.

#### False Discovery Rate Correction

To control for multiple hypothesis testing, we apply a false discovery rate (FDR) correction based on the posterior expected False Discovery Proportion (FDP).

Since differential splicing is evaluated for each junction and factor, we compute factor-specific FDP estimates i.e., to find splicing events enriched or depleted in specific factors. For a given factor *k*, we rank the splice junctions by their posterior probability and estimate the expected FDP at each threshold *τ* as,

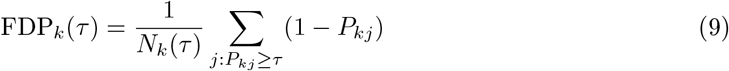

where *N*_*k*_(*τ*) is the number of junctions where *P*_*kj*_ ≥ *τ*. For each factor *k*, the final significance threshold *τ*^*^ is determined as the largest *τ* that satisfies FDP_*k*_(*τ*^*^) ≥ *α*, where *α* (default 0.05) is the FDR control level for factor *k*. This ensures that at most 5% of the reported differentially spliced junctions for a given factor are false positives. This approach follows a Bayesian FDR control strategy similar to scVI but is applied separately for each factor to identify factor-specific differential splicing markers.

### Simulating Junction Usage Ratios Across Cell Types

Our simulations are based on observed ATSE counts and predefined cell types from real TMS brain cells to ensure the overall sparsity of read coverage is maintained. Every 3-junction ATSE is randomly assigned a positive (has DS across cell states) or negative label (no DS) (**Fig. 1A**). The three splice junctions belonging to a given ATSE are sorted into the two corresponding to exon inclusion (**J1** and **J2**) and the one corresponding to exon exclusion (**J3**). For negative ATSEs we sample one shared *ψ*^*J*3^ from Beta(0.5, 0.5) (hyperparameters chosen to emulate the known bimodality of splicing patterns (64)) for all cell states. For positive ATSEs, we sample a different 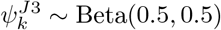 for each cell state *k*. We then set 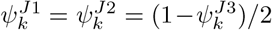 for each *k*. Junction counts are then sampled from binomial distributions with these *ψ* values and total count from the observed (‘real’) TMS data. We evaluate the observed differences, (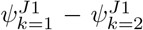 for example) after simulating junction counts to ensure sampled junction usage ratios correspond to their label: positive (DS) ATSE or negative (no DS).

### Modeling Aging with Splicing and Expression

We assessed the predictive power of alternative splicing and gene expression for chronological age using standard linear regression models at both cell-type–specific and global levels. For the cell-type–specific analysis, we iterated over each broad cell type containing at least 50 cells and spanning two or more age groups. Within each group, we trained three separate models to predict numeric age: one using splicing-based latent features (*Φ*), one using gene expression–based NMF features, and one combining both feature sets. We computed *R*^2^ scores using 10-fold cross-validation, examined the relative performance of splicing and expression features, and extracted the top contributing features by absolute model coefficients. A scatter plot of CV *R*^2^ values was used to highlight cell types where either modality offered superior predictive power.

For the global analysis, we trained linear regression models across all cells to compare four combinations of input features: broad cell type alone (one-hot encoded), broad cell type with splicing, broad cell type with expression, and all features combined. Age was modeled as a continuous outcome, and 10-fold cross-validation was used to compute mean and standard deviation of the *R*^2^ score for each model.

### RBP Aging Mediation Analysis

To identify potential regulatory drivers of age-associated splicing changes, we conducted a systematic mediation analysis focusing on a curated list of 269 RNA-binding proteins (RBPs). We hypothesized that chronological age (*X*) influences the expression of specific RBPs (*M*), which in turn modulates the activity of observed SPs (*Y*) (**Fig. 5B**). This analysis was performed independently for each tissue-cell type pair.

We utilized the mediation_analysis function from the pingouin Python package. For each cell, the mediator variable (*M*) was defined as the denoised expression level of a given RBP, obtained from the linear scVI-normalized expression layer to minimize technical noise and dropout effects. The outcome variable (*Y*) was the cell-specific SP activity (*ϕ*) inferred by LeafletFA. The significance of the indirect effect (the mediation effect) was determined using bootstrapping with 500 iterations to estimate 95% confidence intervals and p-values.

To account for multiple hypothesis testing across the suite of RBPs and SPs, we applied the Benjamini-Hochberg False Discovery Rate (FDR) correction to the indirect effect p-values within each cell type. An RBP was considered a significant mediator if the FDR-corrected p-value was less than 0.05. The Aging Mediation Coefficient was defined as the standardized coefficient of the indirect effect, representing the magnitude and direction of the aging signal propagated through RBP expression to the splicing program activity (**Fig. 5C**).

### Cross-species Splicing Conservation Analysis

To evaluate conservation of alternative splicing between human and mouse, we filtered ATSEs in each species to retain only splice junctions derived from genes with one-to-one orthologs, as defined by Ensembl. To focus on simpler splicing topologies, we included only ATSEs involving five or fewer junctions. All annotations were derived from GENCODE vM19 (mouse) and v45 (human), aligned to the GRCm38 (mm10) and GRCh38 (hg38) genome assemblies, respectively.

First, splice junction coordinates were extracted from ATSE definitions and exported to BED format for cross-species comparison. To assess coordinate-level conservation, we used the UCSC liftOver tool to map junction coordinates between assemblies in both directions: mouse junctions were lifted from mm10 to hg38, and human junctions were lifted from hg38 to mm10. The lifted coordinates were then compared against all annotated junctions in the target species to identify possible matches within a ±100 bp window. This proximity-based approach captures biologically equivalent splicing events that may have different coordinates due to evolutionary changes, annotation differences between assemblies, or minor assembly variations.

For sequence-based validation of coordinate matches, we extracted ±50 bp of exonic context flanking donor and acceptor sites from both the lifted junction coordinates and their matched target junction coordinates in the respective genomes. We computed pairwise sequence similarity scores using global alignment and evaluated splice motif conservation (GT–AG, GC–AG). Junctions were classified as *conserved* (≥80% sequence similarity with conserved motifs), *sequence_only* (≥ 80% similarity without motif conservation), *motifs_only* (conserved motifs with <80% similarity), or *diverged* (neither criterion met).

To ensure one-to-one relationships, duplicate mappings were resolved by selecting the junction pair with the highest sequence similarity and lowest coordinate distance. This two-step approach of proximity-based coordinate matching followed by sequence validation identifies high-confidence orthologous splice junctions, ensuring that coordinate proximity represents genuine conservation of the same biological splicing event across species.

## Acknowledgements

We thank Peter Sims, Xuebing Wu and Elham Azizi for helpful feedback. This work was funded by the National Science Foundation (CAREER DBI2146398 to D.A.K.). Any opinions, findings, and conclusions or recommendations expressed in this material are those of the authors and do not necessarily reflect the views of the NSF.

## Author Contributions

K.I. and D.A.K. conceived the study. K.I. developed the LeafletFA framework, performed all data processing, and conducted the primary analysis. D.A.K. supervised the research and provided guidance on model development. K.I. and D.A.K. wrote and edited the manuscript.

## Code Availability

The LeafletFA framework and the ATSEmapper utility are open-source and available on GitHub at https://github.com/daklab/LeafletFA and https://github.com/daklab/LeafletFA-utils. Additionally, all scripts required to reproduce the analyses and figures presented in this manuscript are hosted at https://github.com/karini925/Leaflet-analysis.

## Data Availability

Raw Smart-seq2 data used in this study were obtained from publicly available repositories including the Chan Zuckerberg Biohub (Tabula Muris Senis, Tabula Sapiens) and the Allen Institute for Brain Science. The processed AnnData objects for mouse, including quantified junction counts and inferred SP activities (*Φ*), have been deposited on Zenodo: https://doi.org/10.5281/zenodo.17981150.

## Competing Interests

The authors declare no competing interests.

## Extended Data

### Initializing variational parameters using waypoints

#### Step 1: Initialize ψ_kj_ Using Waypoints

The initialization of *ψ*_*kj*_ is based on the (smoothed) junction usage ratios of a set of selected cells, referred to as waypoints. These cells provide a reference for the factor-specific junction usage, and their observed junction usage ratios are used to initialize 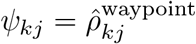.

#### Step 2: Solve for ϕ_ck_

Given *ψ*_*kj*_ and ignoring constraints, we can solve for *ϕ*_*ck*_ by fitting the linear regression model,

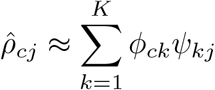

for all cells *c* and junctions *j*. This equation can be solved for *ϕ*_*ck*_ using least squares:

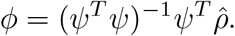

#### Step 3: Bounding and Normalizing ϕ_ck_

To ensure numerical stability and adherence to the constraints of the model, the values of *ϕ*_*ck*_ are clipped to be in [*ϵ*, 1 − *ϵ*],

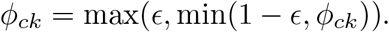

Finally, we normalize *ϕ*_*ck*_ such that the proportions across all factors for each cell sum to 1, reflecting the Dirichlet nature of the cell-factor assignment:

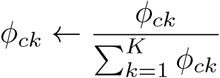

### Evaluating Consistency of Factor Assignments Across Initializations

To assess the reproducibility of factor assignments across different initializations of our probabilistic model, we employed a systematic approach to compare the latent variable assignments. Given the potential for permutation ambiguities in label assignments across different model fits, direct comparison of the cells by factors matrices *ϕ* is non-trivial. Therefore, we used the Hungarian algorithm, which solves bipartite matching problems, to align the factors between pairs of assignment matrices derived from different initializations.

For a pair (*ϕ*^(1)^, *ϕ*^(2)^) (coming from two different initializations) we compute the cost matrix *S* where 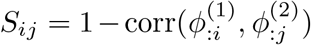 for all *i, j* (indexing factors). The Hungarian algorithm is then used to find the optimal permutation *σ* that minimizes ∑_*i*_ *S*_*iσ*(*i*)_. Factor (column) *i* of *ϕ*^(1)^ is thereby matched with factor (column) *σ*(*i*) of *ϕ*^(2)^.

We calculate the Pearson correlation coefficient for each matched pair of factors post ‘alignment’ and the overall model stability or consistency is summarized by calculating the median of these coefficients across initialization pairs.

**Supplementary Figure 1:**
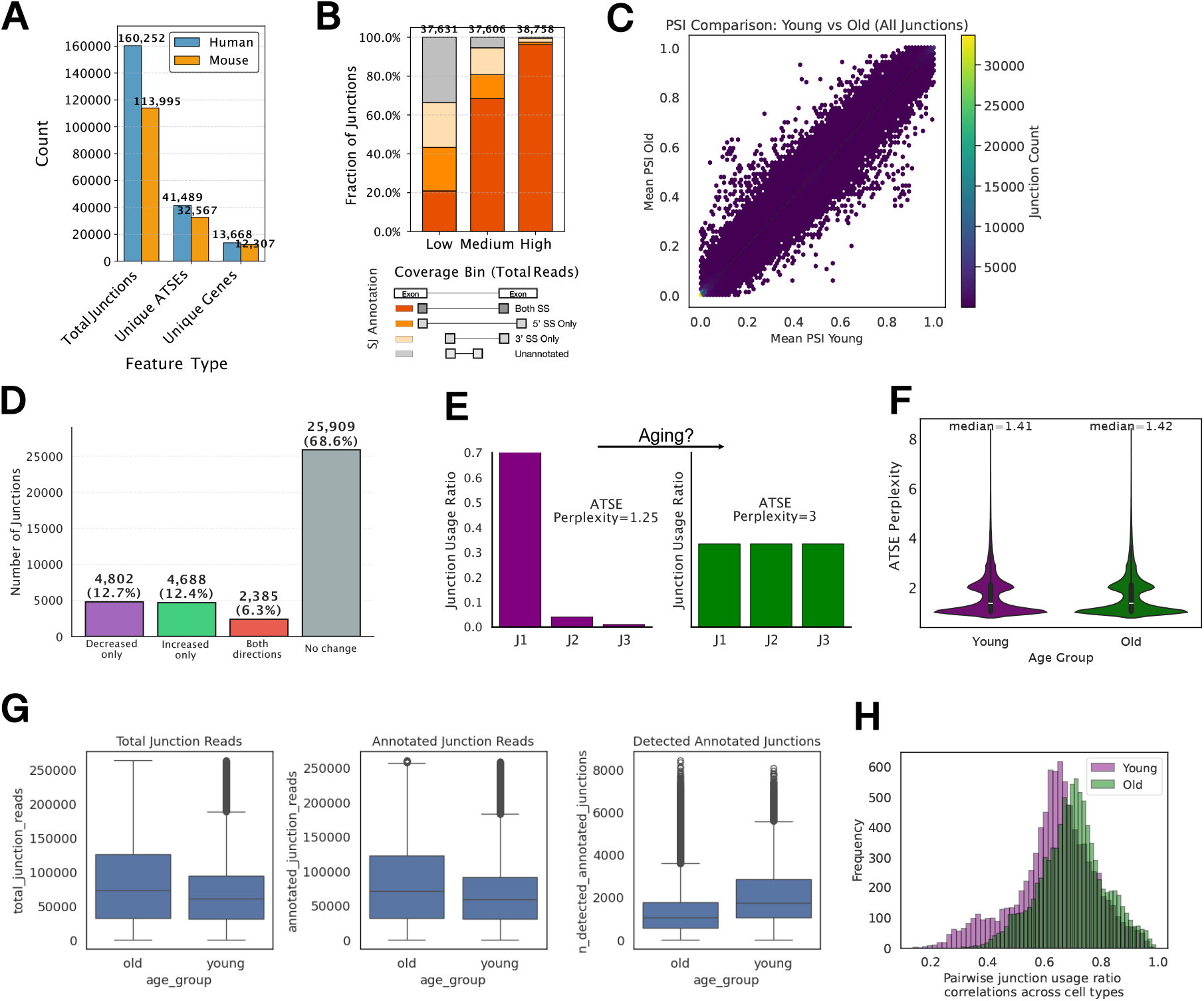
Overview of ATSEmapper results in mouse and human Alternative Splicing aging variation. **(A)** Simulation workflow schematic. 1) Identify exon skipping ATSEs with three junctions (J1, J2, J3). 2) Randomly assign each ATSE as positive (DS) or negative (no DS). 3) Simulate junction usage ratios (*Ψ*) using Beta distribution: if ATSE is positive, allow junction *Ψ* to vary across K cell groups; if negative, no significant variation across K. 4) Simulate junction counts from observed cell ATSE counts and simulated PSI using Binomial distribution. **(B)** Silhouette score comparison between LeafletFA and NMF across varying percentages of DS (differentially spliced) junctions (10%, 50%, 90%). **(C)** Silhouette scores by percentage of positive DS events and number of latent factors K (K=2 vs K=17). **(D)** Spearman correlation between simulated and estimated delta values in junction usage ratios across factors, comparing Beta-Binomial and Binomial likelihoods across different percentages of positive DS events. **(E)** Precision analysis comparing junction-specific shape prior (YES vs NO) and likelihood models (Beta-Binomial vs Binomial) for simulations with 10% DS events (left) and >10% DS events (right). **(F)** Precision-recall curve for LeafletFA with area under curve (AP = 0.94) showing model performance in detecting differentially spliced events. **(G)** Silhouette scores for simulations with >10% DS events, comparing junction-specific shape prior (YES vs NO) and likelihood models (Beta-Binomial vs Binomial). **(H)** UMAP visualization of *Φ* (cell factor activity) embeddings for the model configuration achieving the highest silhouette score (0.82) on simulated data, showing 17 distinct cell states colored by simulated cell group assignment.

**Supplementary Figure 2:**
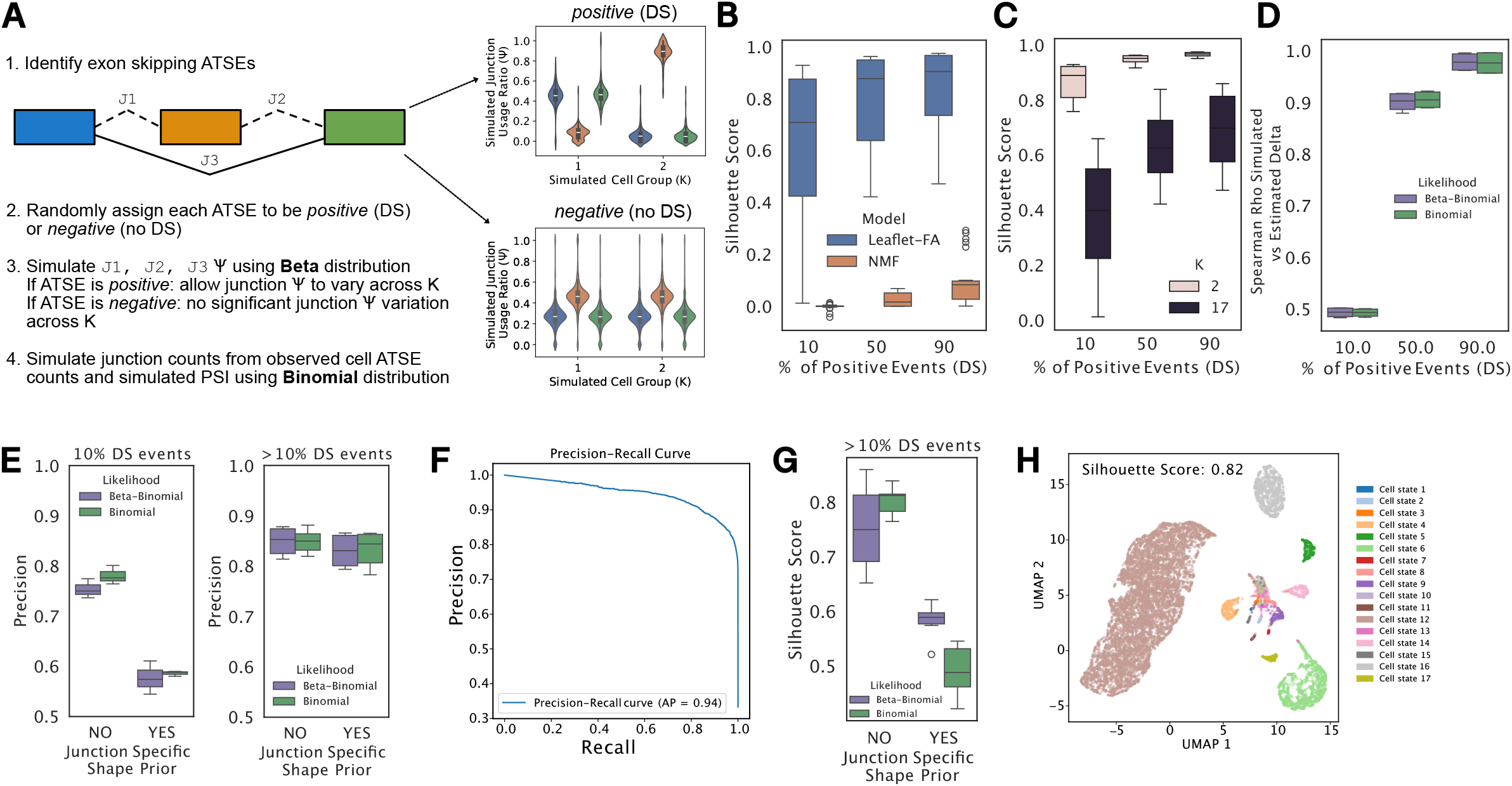
LeafletFA performance evaluation on simulated single-cell junction count data. **(A)** Simulation workflow schematic. 1) Identify exon skipping ATSEs with three junctions (J1, J2, J3). 2) Randomly assign each ATSE as positive (DS) or negative (no DS). 3) Simulate junction usage ratios (*Ψ*) using Beta distribution: if ATSE is positive, allow junction *Ψ* to vary across K cell groups; if negative, no significant variation across K. 4) Simulate junction counts from observed cell ATSE counts and simulated PSI using Binomial distribution. **(B)** Silhouette score comparison between LeafletFA and NMF across varying percentages of DS (differentially spliced) junctions (10%, 50%, 90%). **(C)** Silhouette scores by percentage of positive DS events and number of latent factors K (K=2 vs K=17). **(D)** Spearman correlation between simulated and estimated delta values in junction usage ratios across factors, comparing Beta-Binomial and Binomial likelihoods across different percentages of positive DS events. **(E)** Precision analysis comparing junction-specific shape prior (YES vs NO) and likelihood models (Beta-Binomial vs Binomial) for simulations with 10% DS events (left) and >10% DS events (right). **(F)** Precision-recall curve for LeafletFA with area under curve (AP = 0.94) showing model performance in detecting differentially spliced events. **(G)** Silhouette scores for simulations with >10% DS events, comparing junction-specific shape prior (YES vs NO) and likelihood models (Beta-Binomial vs Binomial). **(H)** UMAP visualization of *Φ* (cell factor activity) embeddings for the model configuration achieving the highest silhouette score (0.82) on simulated data, showing 17 distinct cell states colored by simulated cell group assignment.

**Supplementary Figure 3:**
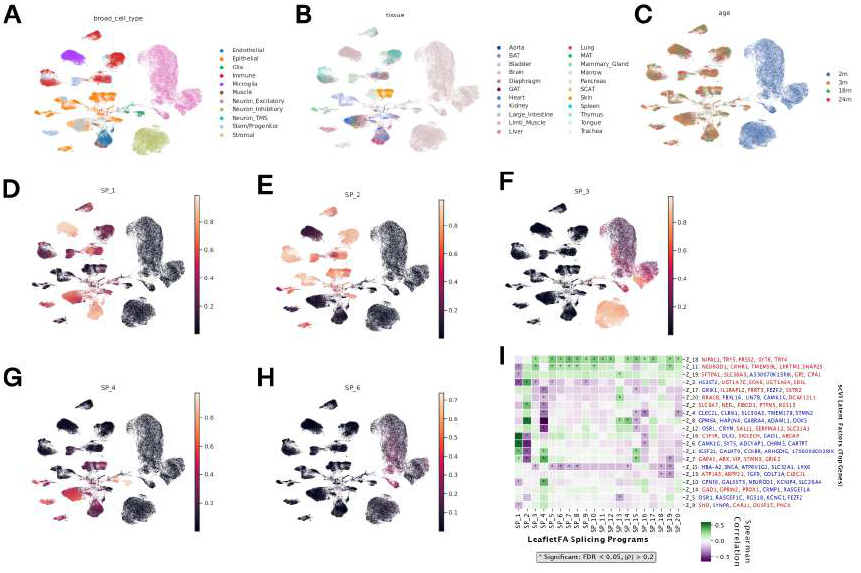
-cell landscape of splicing programs and cell-type heterogeneity in the Smart-seq2 mouse atlas. **(A-C)** UMAP visualization of all cells across all cell types in the Smart-seq2 mouse atlas. UMAP embeddings were computed based on scVI latent factors derived from gene expression. Cells are color-coded by **(A)** broad cell type, **(B)** tis,;ue of origin, and **(C)** animal age (2, 3, 18, and 24 months). **(D-H)** Spatial distributio11 of LeafletFA Splicing Program (SP) activity scores projected onto the gene expression-based Ul’v1AP manifold for representative programs **(D)** SP_ 1, (E) SP_ 2, **(F)** SP_ 3, (**G)** SP_ 4, and **(H)** SP_ 6. Color scales represent normalized activity levels, illustrating the cell-type and tissue specificity of distinct splicing signatures. **(I)** Correlation matrix between LeafletFA SPs and scVI gene expression latent factors *(Z* = 20) with hierarchical clustering performed on *Z*. Asterisks indicate significant Spearman correlations (FDR < 0.05, | *ρ* | > 0.2). Top genes associated with each scVI latent factor are listed on the right; gene names in red indicate positive loadings in *Z*, while those in blue indicate negative loadings.

**Supplementary Figure 4:**
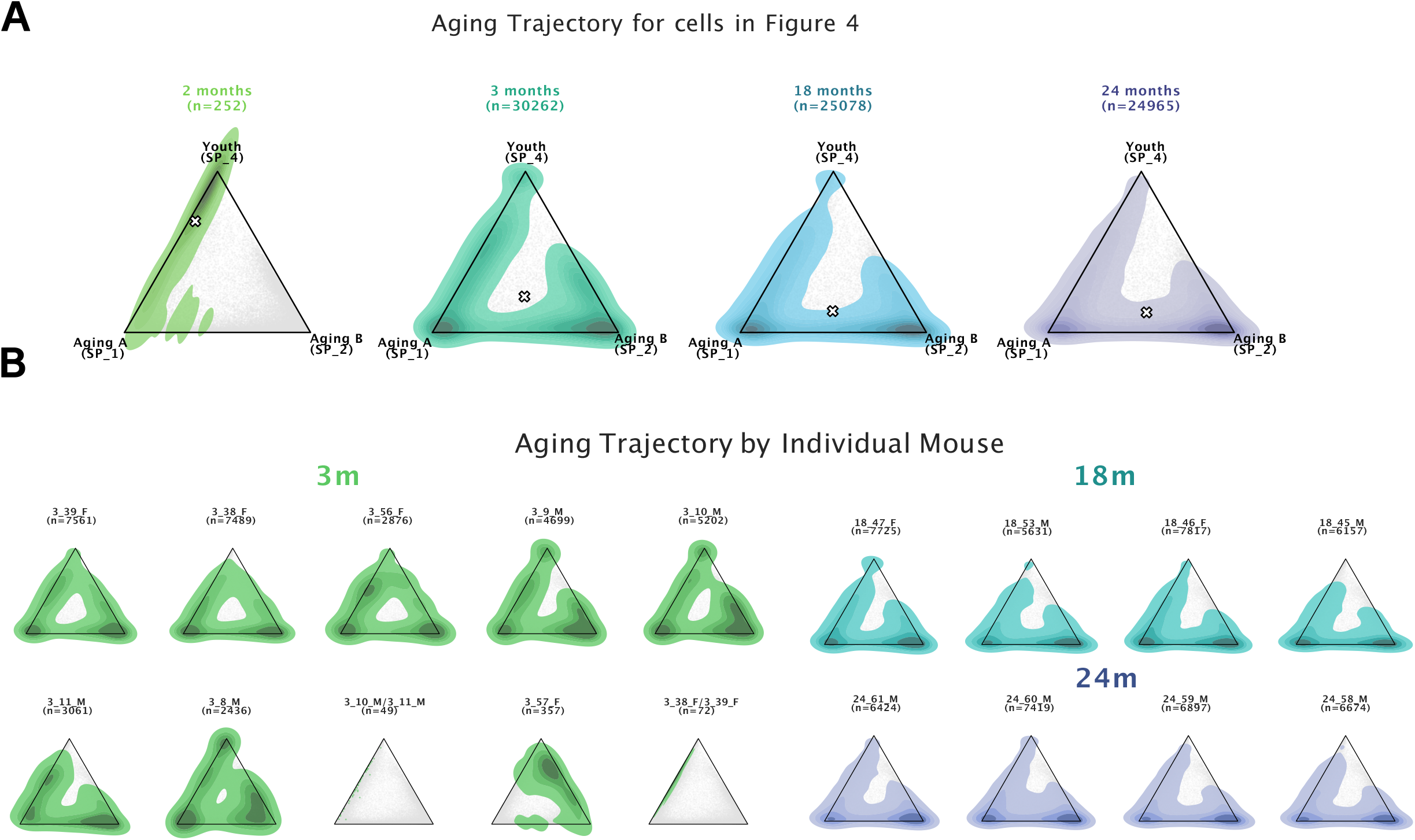
Robustness and Individual Consistency of Structural Aging Trajectories. **(A)** Faceted ternary plots show the distribution of splice-program (SP) activity for the 76 significant cell types identified in Figure 4 across four age groups (2, 3, 18, and 24 months). Each plot represents the relative activity of Youth (SP4, top), Aging A (SP1, left), and Aging B (SP2, right). The colored contours represent the Kernel Density Estimation (KDE) of cell density for that age group, while the light grey background indicates the total landscape of all cells from these 76 cell types across all ages. The white ‘x’ indicates the mean coordinate (center of mass) for each age group. **(B)** Ternary density plots faceted by individual mouse and age demonstrate the robustness of the aging shift. This analysis is restricted to mice from the TMS dataset because the AB dataset only provides 2-month-old nuclei, making it unsuitable for individual aging trajectory comparisons. Each subplot shows the KDE for a single donor. Sample sizes (*n*) for each individual are indicated above each plot.

**Supplementary Figure 5:**
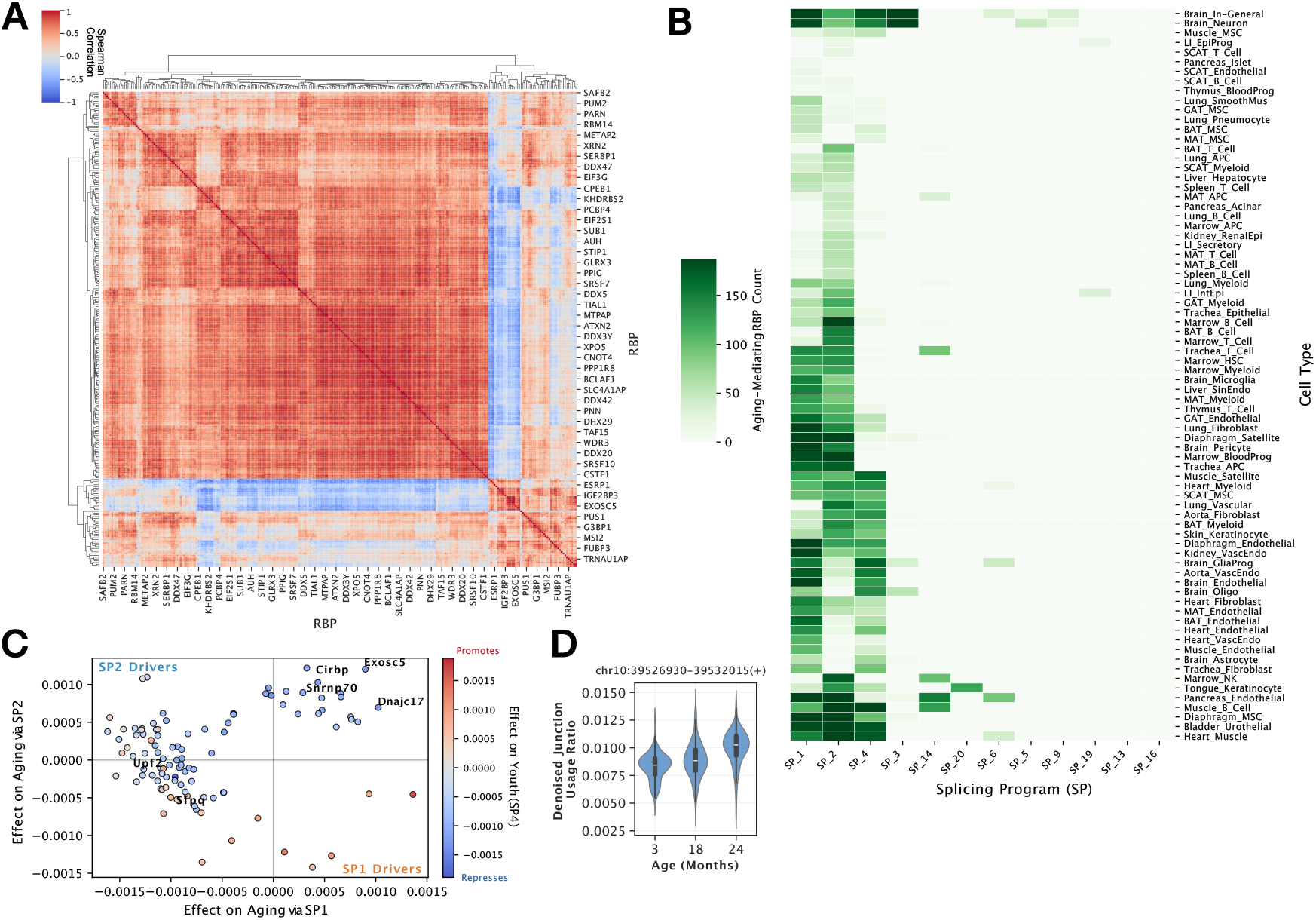
Systematic identification and global landscape of aging-associated RBP mediators. **(A)** RBP co-expression and regulatory modules. Spearman correlation heatmap illustrating the expression relationships between identified RNA-binding protein (RBP) candidates across all tissues and cell types. The hierarchical clustering identifies distinct modules of coregulated splicing factors, suggesting that aging-associated splicing shifts may be governed by redundant or coordinated regulatory units rather than single drivers. **(B)** Cell-type specificity of aging mediators. Heatmap showing the count of significant aging-mediating RBPs identified across diverse tissues and cell types (y-axis) for individual Splicing Programs (x-axis). **(C)** Divergent mediation effects on youth vs. aging programs. Scatter plot correlating RBP mediation coefficients for SP1 (x-axis) and SP2 (aging; y-axis). Data points are colored by their effect on the youth program (SP4), revealing that top aging drivers like *Cirbp, Exosc5*, and *Snrnp70* consistently promote aging-associated programs while repressing the youth program. **(D)** Minimal usage of the *Fyn* green junction. Violin plots showing the distribution of denoised junction usage ratios (imputed PSI) for the specific exon-skipping junction in *Fyn* across age groups. This junction (represented by green in Figure 5F) is utilized very little across the lifespan in bladder urothelial cells.

**Supplementary Figure 6:**
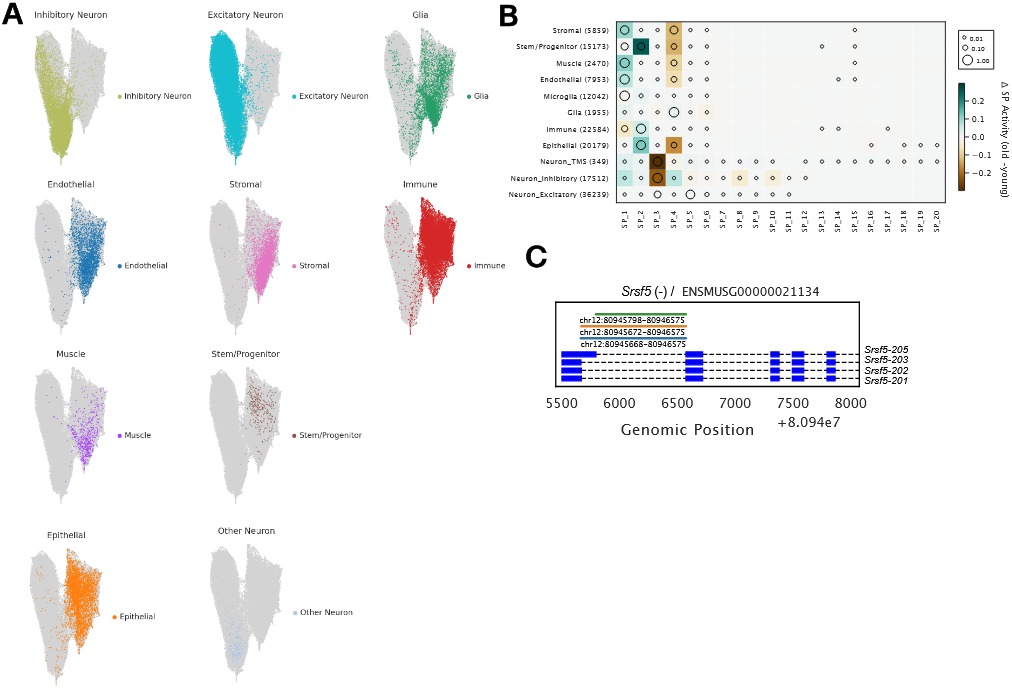
Cross-species transfer learning validation and conserved *SRSF5* splicing dynamics. **(A)** Conserved cell identity recovery via transfer learning. UMAP visualization of human cells where coordinates are derived solely from the activities of orthologous mouse splicing programs (*Φ*_*H*_). Cells are colored by their broad cell type, demonstrating that conserved splicing signatures are sufficient to recover discrete biological identities across species. **(B)** Global age-associated SP activity shifts in mouse. Dot plot illustrating the change in SP activity between young and old cohorts across major broad mouse cell types. The color indicates the direction and magnitude of the shift (teal for increase, brown for decrease), while dot size represents the median SP activity within the lineage. **(C)** Conserved *Srsf5* alternative transcription and splicing event in mouse. Genomic track view of the murine *Srsf5* locus (ENSMUSG00000021134). Similar to the human ortholog, the locus features alternative 5’ SS that differ by four nucleotides.

## Notes

### Competing Interest Statement

The authors have declared no competing interest.

### Summary of Updates

LaTeX-specific syntax was removed from the abstract.

## Bibliography

[1] C. López-Otín et al. The hallmarks of aging. Cell, 153(6):1194–1217, June 2013.

[2] F. E. Baralle and J. Giudice. Alternative splicing as a regulator of development and tissue identity. Nat. Rev. Mol. Cell Biol., 18(7):437–451, July 2017.

[3] C. Heintz et al. Splicing factor 1 modulates dietary restriction and TORC1 pathway longevity in C. elegans. Nature, 541(7635):102–106, January 2017.

[4] C. M.-C. Li et al. Aging-associated alterations in mammary epithelia and stroma revealed by single-cell RNA sequencing. Cell Rep., 33(13):108566, December 2020.

[5] K. Jin et al. Brain-wide cell-type-specific transcriptomic signatures of healthy ageing in mice. Nature, pp. 1–15, January 2025.

[6] K. Rhine et al. Neuronal aging causes mislocalization of splicing proteins and unchecked cellular stress. Nat. Neurosci., 28(6):1174–1184, June 2025.

[7] S. A. Rodríguez et al. Global genome splicing analysis reveals an increased number of alternatively spliced genes with aging. Aging Cell, 15(2):267–278, April 2016.

[8] L. W. Harries et al. Human aging is characterized by focused changes in gene expression and deregulation of alternative splicing: Gene expression changes in human aging. Aging Cell, 10(5):868–878, October 2011.

[9] S. A. Rodríguez et al. Global genome splicing analysis reveals an increased number of alternatively spliced genes with aging. Aging Cell, 15(2):267–278, April 2016.

[10] M. Mariotti et al. Deterioration of the human transcriptome with age due to increasing intron retention and spurious splicing. bioRxiv, pp. 2022.03.14.484341, March 2022.

[11] Y. Song et al. Single-cell alternative splicing analysis with expedition reveals splicing dynamics during neuron differentiation. Mol. Cell, 67(1):148–161.e5, July 2017.

[12] D. Lukacsovich et al. Single-cell RNA-seq reveals developmental origins and ontogenetic stability of neurexin alternative splicing profiles. Cell Rep., 27(13):3752–3759.e4, June 2019.

[13] A. S. Booeshaghi et al. Isoform cell-type specificity in the mouse primary motor cortex. Nature, 598(7879):195–199, October 2021.

[14] G. Benegas et al. Robust and annotation-free analysis of alternative splicing across diverse cell types in mice. Elife, 11, March 2022.

[15] Tabula Muris Consortium. A single-cell transcriptomic atlas characterizes ageing tissues in the mouse. Nature, 583(7817):590–595, July 2020.

[16] Tabula Sapiens Consortium* et al. The tabula sapiens: A multiple-organ, single-cell transcriptomic atlas of humans. Science, 376(6594):eabl4896, May 2022.

[17] R. D. Hodge et al. Conserved cell types with divergent features in human versus mouse cortex. Nature, 573(7772):61–68, September 2019.

[18] Tabula Sapiens Consortium* et al. The tabula sapiens: A multiple-organ, single-cell transcriptomic atlas of humans. Science, 376(6594):eabl4896, May 2022.

[19] J. E. Olivieri et al. The SpliZ generalizes ‘percent spliced in’ to reveal regulated splicing at single-cell resolution. Nat. Methods, 19(3):307–310, March 2022.

[20] C. F. Buen Abad Najar et al. Identifying cell state–associated alternative splicing events and their coregulation. Genome Res., July 2022.

[21] S. M. Weyn-Vanhentenryck et al. Precise temporal regulation of alternative splicing during neural development. Nat. Commun., 9(1):2189, June 2018.

[22] D. F. Moakley et al. Reverse engineering neuron-type-specific and type-orthogonal splicing-regulatory networks using diverse cellular transcriptomes. Cell Rep., 44(7):115898, June 2025.

[23] H. Feng et al. Complexity and graded regulation of neuronal cell-type-specific alternative splicing revealed by single-cell RNA sequencing. Proc. Natl. Acad. Sci. U. S. A., 118(10), March 2021.

[24] M. Jacko et al. Rbfox splicing factors promote neuronal maturation and axon initial segment assembly. Neuron, 97(4):853–868.e6, February 2018.

[25] C. C. Warzecha et al. An ESRP-regulated splicing programme is abrogated during the epithelial-mesenchymal transition. EMBO J., 29(19):3286–3300, October 2010.

[26] R. Z. Kunes et al. Supervised discovery of interpretable gene programs from single-cell data. Nat. Biotechnol., 42(7):1084–1095, July 2024.

[27] K. C. Cotto et al. RegTools: Integrated analysis of genomic and transcriptomic data for the discovery of splicing variants in cancer. April 2021.

[28] M. D. Schertzer et al. Perplexity as a metric for isoform diversity in the human transcriptome. bioRxiv, pp. 2025.07.02.662769, July 2025.

[29] F. Reese et al. The ENCODE4 long-read RNA-seq collection reveals distinct classes of transcript structure diversity. bioRxiv, pp. 2023.05.15.540865, May 2023.

[30] Y. I. Li et al. Annotation-free quantification of RNA splicing using LeafCutter. Nat. Genet., 50(1):151–158, January 2018.

[31] B. Eli et al. Pyro: Deep universal probabilistic programming. arXiv [cs.LG], October 2018.

[32] C. C. Warzecha et al. ESRP1 and ESRP2 are epithelial cell-type-specific regulators of FGFR2 splicing. Mol. Cell, 33(5):591–601, March 2009.

[33] S. Bocklandt et al. Epigenetic predictor of age. PLoS One, 6(6):e14821, June 2011.

[34] S. Horvath. DNA methylation age of human tissues and cell types. Genome Biol., 14(10):R115, December 2013.

[35] J.-H. Yang et al. Loss of epigenetic information as a cause of mammalian aging. Cell, January 2023.

[36] M. A. Argentieri et al. Proteomic aging clock predicts mortality and risk of common agerelated diseases in diverse populations. Nat. Med., 30(9):2450–2460, September 2024.

[37] K. Jin et al. Brain-wide cell-type-specific transcriptomic signatures of healthy ageing in mice. Nature, pp. 1–15, January 2025.

[38] M. J. Zhang et al. Mouse aging cell atlas analysis reveals global and cell type-specific aging signatures. Elife, 10, April 2021.

[39] E. Rosonina and B. J. Blencowe. Gene expression: the close coupling of transcription and splicing. Curr. Biol., 12(9):R319–21, April 2002.

[40] D. L. Bentley. Coupling mRNA processing with transcription in time and space. Nat. Rev. Genet., 15(3):163–175, March 2014.

[41] S. Loerch et al. The pre-mRNA splicing and transcription factor tat-SF1 is a functional partner of the spliceosome SF3b1 subunit via a U2AF homology motif interface. J. Biol. Chem., 294(8):2892–2902, February 2019.

[42] U. Braunschweig et al. Dynamic integration of splicing within gene regulatory pathways. Cell, 152(6):1252–1269, March 2013.

[43] H. E. Merens et al. Timing is everything: advances in quantifying splicing kinetics. Trends Cell Biol., 0(0), May 2024.

[44] S. Naftelberg et al. Regulation of alternative splicing through coupling with transcription and chromatin structure. Annu. Rev. Biochem., 84:165–198, 2015.

[45] M. Deschênes and B. Chabot. The emerging role of alternative splicing in senescence and aging. Aging Cell, 16(5):918–933, October 2017.

[46] M. Franck et al. Nonuniversality of inflammaging across human populations. Nat. Aging, pp. 1–10, June 2025.

[47] N. Thrupp et al. Single-nucleus RNA-seq is not suitable for detection of microglial activation genes in humans. Cell Rep., 32(13):108189, September 2020.

[48] B. Fair et al. Global impact of unproductive splicing on human gene expression. Nat. Genet., 56(9):1851–1861, September 2024.

[49] J. T. Mendell et al. Nonsense surveillance regulates expression of diverse classes of mammalian transcripts and mutes genomic noise. Nat. Genet., 36(10):1073–1078, October 2004.

[50] C. Mertes et al. Detection of aberrant splicing events in RNA-seq data using FRASER. Nat. Commun., 12(1):529, January 2021.

[51] S. Vaidyanathan et al. Robust integration of sparse single-cell alternative splicing and gene expression data with SpliceVI. bioRxiv, pp. 2025.11.26.690853, December 2025.

[52] T. Ashuach et al. MultiVI: deep generative model for the integration of multimodal data. Nat. Methods, 20(8):1222–1231, August 2023.

[53] A. A. Moinfar and F. J. Theis. Unsupervised deep disentangled representation of single-cell omics. bioRxiv, pp. 2024.11.06.622266, November 2024.

[54] Z. Wolfe et al. Deep transcriptomics reveals cell-specific isoforms of pan-neuronal genes. Nat. Commun., 16(1):4507, May 2025.

[55] Z.-X. Shi et al. High-throughput and high-accuracy single-cell RNA isoform analysis using PacBio circular consensus sequencing. Nat. Commun., 14(1):2631, May 2023.

[56] A. B. Rosenberg et al. Single-cell profiling of the developing mouse brain and spinal cord with split-pool barcoding. Science, 360(6385):176–182, April 2018.

[57] A. Sziraki et al. A global view of aging and alzheimer’s pathogenesis-associated cell population dynamics and molecular signatures in human and mouse brains. Nat. Genet., 55(12):2104– 2116, December 2023.

[58] Z. Yao et al. A taxonomy of transcriptomic cell types across the isocortex and hippocampal formation. Cell, 184(12):3222–3241.e26, June 2021.

[59] S. R. Quake and The Tabula Sapiens Consortium. Tabula sapiens reveals transcription factor expression, senescence effects, and sex-specific features in cell types from 28 human organs and tissues. bioRxiv, pp. 2024.12.03.626516, December 2024.

[60] B. Phipson et al. Gene length and detection bias in single cell RNA sequencing protocols. F1000Res., 6:595, April 2017.

[61] V. Svensson et al. Interpretable factor models of single-cell RNA-seq via variational autoencoders. Bioinformatics, 36(11):3418–3421, June 2020.

[62] I. Virshup et al. anndata: Access and store annotated data matrices. J. Open Source Softw., 9(101):4371, September 2024.

[63] M. Setty et al. Characterization of cell fate probabilities in single-cell data with palantir. Nat. Biotechnol., 37(4):451–460, April 2019.

[64] C. F. Buen Abad Najar et al. Coverage-dependent bias creates the appearance of binary splicing in single cells. Elife, 9, June 2020.

